# Spatially-Resolved Multiomic Atlas of Leiomyosarcoma Identifies Two Clinically Relevant Epigenetically-Driven Cell States

**DOI:** 10.64898/2026.05.22.726988

**Authors:** Ryan A. Denu, Zhao Zheng, Yingda Jiang, Suresh Satpati, Veena Kochat, Kumaren Anand, Danh D. Truong, Andrew R. Lynch, William Padron, Davis R. Ingram, Khalida M. Wani, Diana Shamsutdinova, Angela D. Bhalla, Sharon M. Landers, Ravin Ratan, Neeta Somaiah, Christina L. Roland, Keila E. Torres, Peter Van Loo, Pamela T. Soliman, Larissa A. Meyer, Alexander J. Lazar, Emily Z. Keung, Elise F. Nassif Haddad, Kunal Rai

**Author notes:** Co-corresponding authors **Corresponding authors:** Kunal Rai, The University of Texas MD Anderson Cancer Center, Houston, TX 77030., Elise Nassif Haddad, The University of Texas MD Anderson Cancer Center, Houston, TX 77030., Emily Keung, The University of Texas MD Anderson Cancer Center, Houston, TX 77030.

## Abstract

Leiomyosarcoma is a smooth muscle–derived malignancy marked by significant clinical heterogeneity. The extent and nature of cellular heterogeneity and molecular underpinnings remain poorly understood. To address this at transcriptomic and epigenomic levels, we performed single-nucleus multiome sequencing on untreated primary leiomyosarcoma tissues. Malignant cells segregated almost exclusively into two previously unrecognized and epigenetically distinct states: a dedifferentiated, mesenchymal-like subtype (MES) and a differentiated smooth muscle–enriched subtype (SMC). Chromatin accessibility profiling revealed strong enrichment of nuclear factor I (NFI) transcription factor motifs in MES cells, whereas AP-1 family motifs—most prominently FOSL2—were selectively accessible in SMC cells. Established leiomyosarcoma cell lines faithfully recapitulated these subtypes, and targeted depletion of NFI or AP-1 factors suppressed proliferation, invasion, and *in vivo* tumor growth, demonstrating functional dependency on these transcriptional programs. Spatial transcriptomics across 328 tissue cores from 128 leiomyosarcomas showed that immunosuppressive macrophages preferentially cluster around MES regions, revealing a subtype-specific tumor–immune niche. Clinically, MES-dominant tumors were associated with significantly worse patient outcomes. Through an epigenetic inhibitor screen, we identify and validate SMARCA4/2 inhibition as a promising therapeutic vulnerability for MES leiomyosarcomas. Together, this work defines two epigenetically driven, transcription factor–regulated, and clinically relevant states of leiomyosarcoma, revealing mechanistic underpinnings of tumor heterogeneity and uncovering actionable therapeutic strategies.

**GRAPHICAL ABSTRACT:** 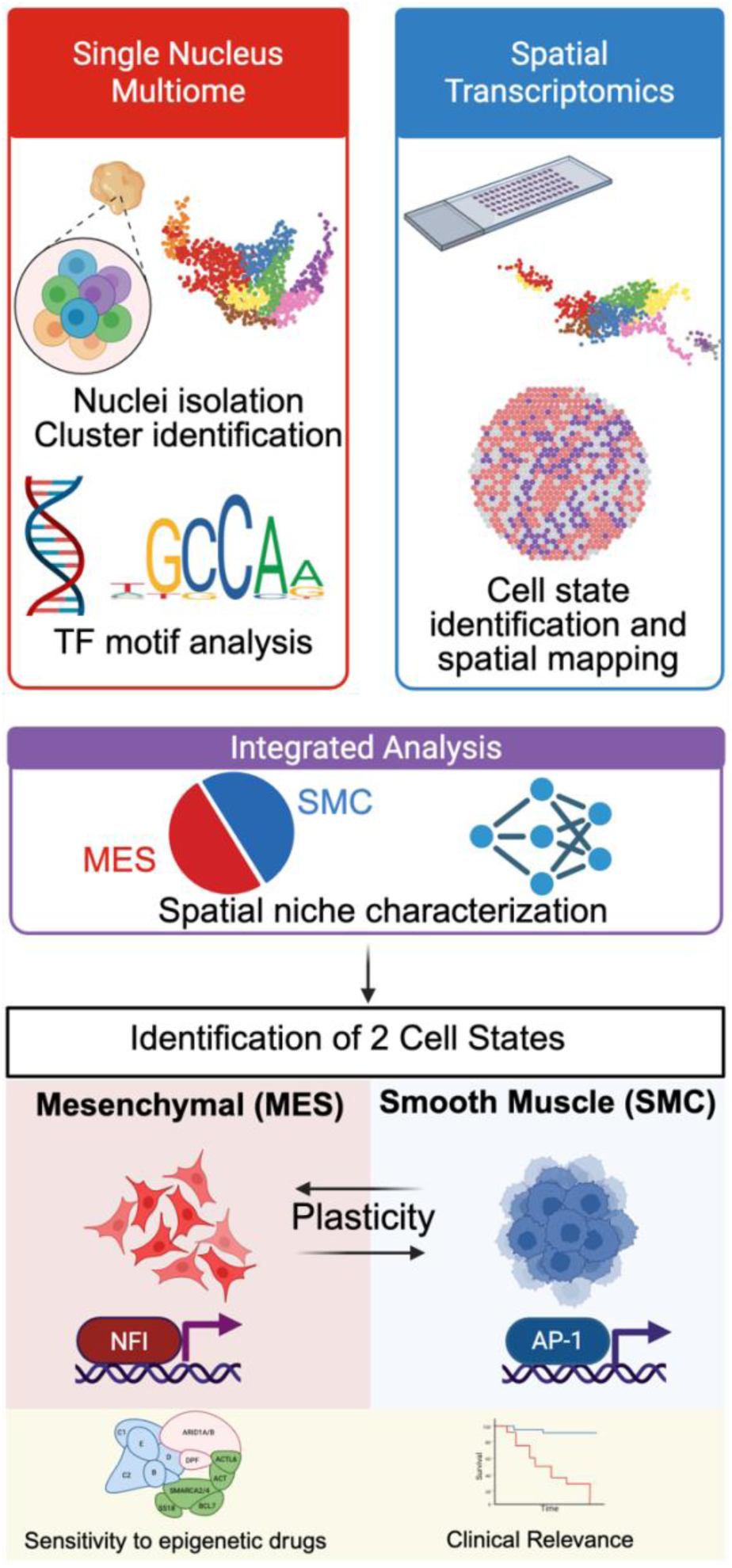

## INTRODUCTION

Leiomyosarcoma is one of the most common subtypes of soft tissue sarcoma (∼20%) and originates from smooth muscle cells, most frequently in the retroperitoneum and uterus. These tumors are aggressive and have high rates of recurrence. Treatment for recurrent or metastatic leiomyosarcoma typically relies on combination cytotoxic chemotherapy, yet response rates remain only 20–30%, and median survival is approximately 30 months.^1^ Consequently, there is a critical need to better understand leiomyosarcoma biology and to develop more effective therapies.

Leiomyosarcoma is a heterogeneous disease with diverse clinical behavior, histologic subtypes, and range of responses to chemotherapy. Previous studies have attempted to characterize this heterogeneity and define its genomic and transcriptomic drivers. These efforts consistently identified three transcriptomic subtypes.^2–8^ However, the translation of these findings into clinical practice has been hampered by substantial inconsistency in the gene sets used to define subtypes across studies. In addition, these analyses relied on bulk sequencing, which obscures the contributions of individual cell types and states. Recently developed single-cell multiomics approaches—which have not yet been applied to leiomyosarcoma—offer a powerful means of characterizing transcriptional, epigenomic, and proteomic heterogeneity at single-cell resolution. Integrative analyses of such datasets have proven highly effective in identifying therapeutic vulnerabilities, exemplified by the discovery of BCL2 as a target to overcome asparaginase resistance in B-cell acute lymphoblastic leukemia.^9^

Although genomic insights have led to transformative targeted therapies in many cancers, this has not been the case for leiomyosarcoma, in which clinically actionable alterations are uncommon. The most frequently mutated genes—*TP53*, *ATRX*, *RB1*, and *MED12*^2,10–12^—currently offer limited therapeutic opportunities. Moreover, the low prevalence of clear driver mutations and the characteristically low tumor mutational burden suggest that leiomyosarcoma oncogenesis and heterogeneity may be driven primarily by non-genetic mechanisms. Epigenetic regulation is now recognized as a hallmark of cancer and plays essential roles in differentiation, proliferation, and genomic stability. Notably, two of the most commonly altered genes in leiomyosarcoma, *ATRX* and *MED12*, are directly involved in epigenetic reprogramming. We therefore hypothesized that epigenomic remodeling underlies leiomyosarcoma heterogeneity and defines subtypes with distinct clinical behaviors.

To date, the epigenomic landscape of leiomyosarcoma has remained largely unexplored. The aim of this study was to delineate the epigenetic regulatory programs that drive leiomyosarcoma heterogeneity at single-cell and spatial resolution. Using single-nucleus multiome sequencing of primary human tumors, we identified two previously unrecognized cellular subtypes. We further applied custom spatial transcriptomic analysis to characterize the distinct microenvironmental niches associated with each subtype. Finally, we leveraged *in vitro* and *in vivo* patient-derived leiomyosarcoma models to uncover a novel therapeutic vulnerability within this newly defined framework.

## RESULTS

### Single nucleus atlas of leiomyosarcoma

To characterize cellular and molecular heterogeneity in leiomyosarcoma, we performed single-nucleus multiome sequencing to jointly profile gene expression (snRNA-seq) and chromatin accessibility (snATAC-seq). We analyzed 16 untreated primary leiomyosarcoma samples derived from three anatomic sites: retroperitoneal (n = 9), extremity (n = 3), and uterus (n = 4) (**Figure 1A**). Tumor identity and purity were confirmed by two independent expert sarcoma pathologists (**Supplementary Figure 1**). Clinical characteristics are summarized in **Table 1** and are notable for a mix of disease stages and mostly high-grade tumors (13/16), although some intermediate-grade (3/16) tumors were included to capture leiomyosarcoma heterogeneity. Low-grade tumors were excluded due to their distinct biology and clinical outcomes relative to intermediate- and high-grade leiomyosarcoma. In total, we obtained high quality snRNA-seq data from 94,439 cells (**Supplementary Table S1**). UMAP analysis identified 15 distinct cell clusters (**Figure 1B**).

**Figure 1.**
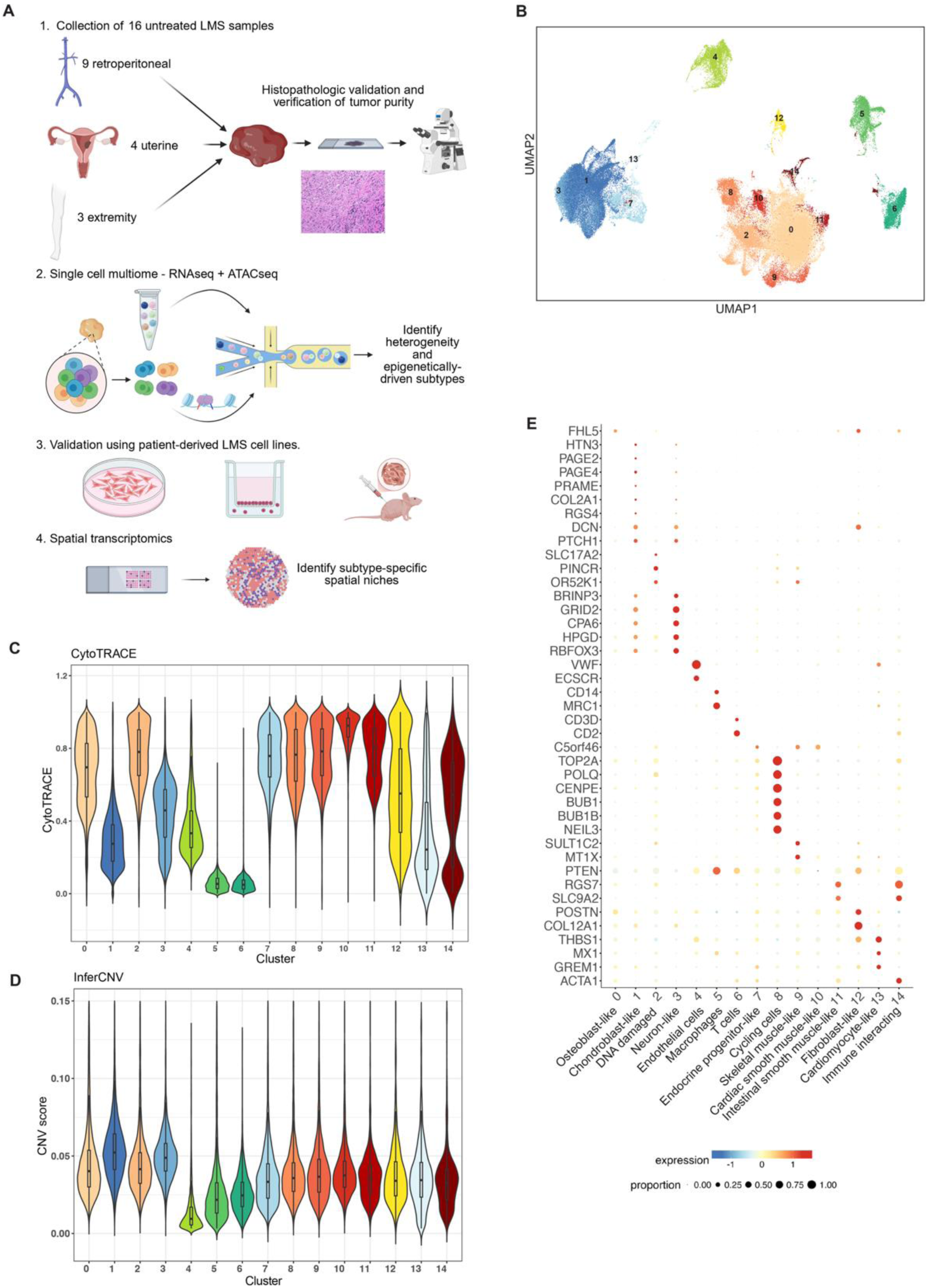
Single cell atlas of leiomyosarcoma. (A) Study schema. (B) snRNA-seq UMAP analysis of all 16 leiomyosarcoma samples. (C) Plot of CytoTRACE scores by cluster. (D) Plot of InferCNV scores by cluster. (E) Bubble plot of a subset of the most differentially expressed genes (DEGs) for each cluster. Annotations are shown on the bottom based on pathway analysis and annotation of the DEGs.

**Table 1.**
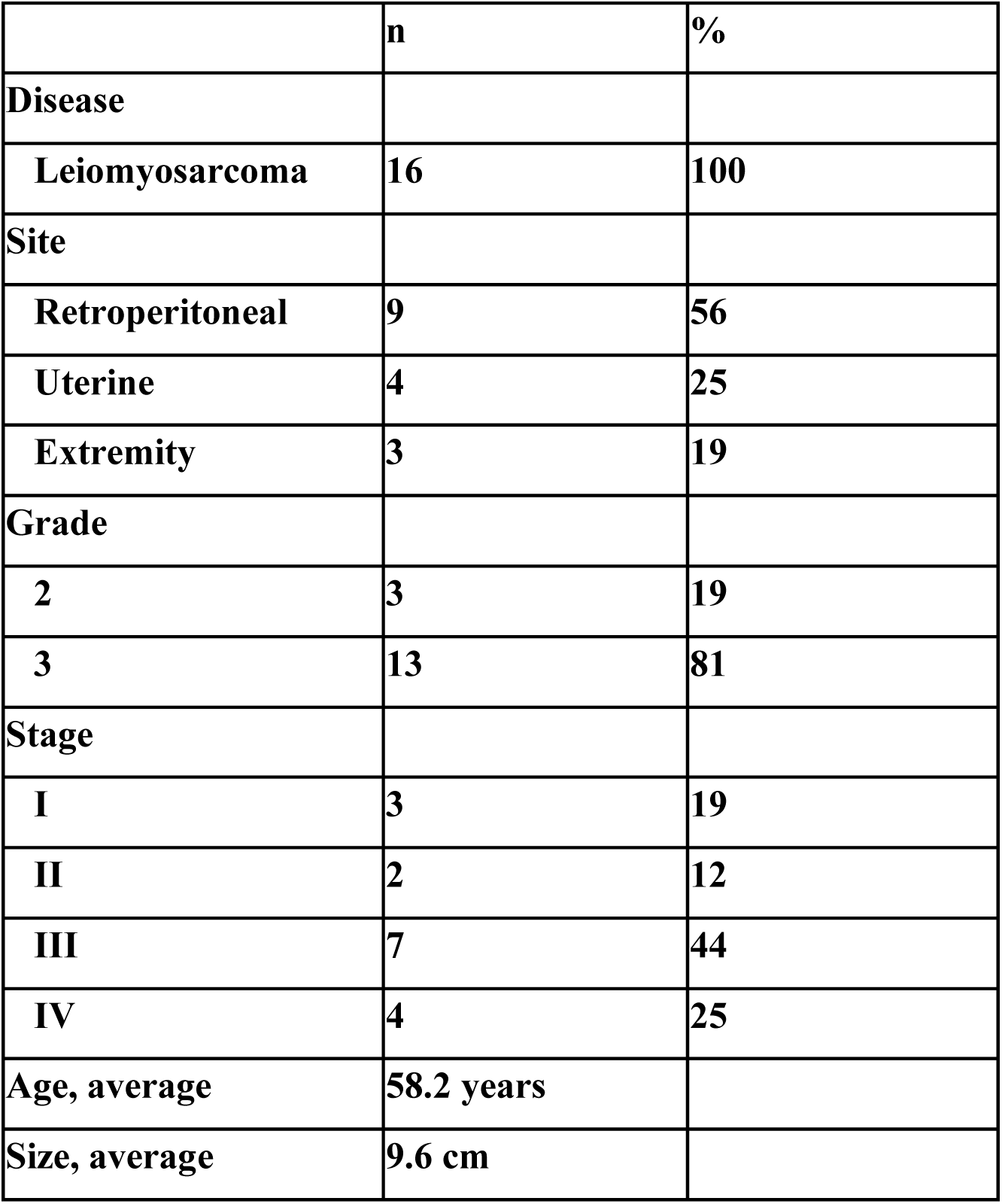
Patient characteristics.

|  | **n** | **%** |
| --- | --- | --- |
| **Disease** |  |  |
| **Leiomyosarcoma** | **16** | **100** |
| **Site** |  |  |
| **Retroperitoneal** | **9** | **56** |
| **Uterine** | **4** | **25** |
| **Extremity** | **3** | **19** |
| **Grade** |  |  |
| **2** | **3** | **19** |
| **3** | **13** | **81** |
| **Stage** |  |  |
| **I** | **3** | **19** |
| **II** | **2** | **12** |
| **III** | **7** | **44** |
| **IV** | **4** | **25** |
| **Age, average** | **58.2 years** |  |
| **Size, average** | **9.6 cm** |  |

To distinguish benign from malignant cells, we applied two complimentary approaches. First, we used CytoTRACE, which infers differentiation state based on the number of genes expressed per cell.^13^ Clusters 4–6 exhibited lower CytoTRACE scores (**Figure 1C**). Second, we assessed copy-number variation (CNV) using InferCNV (**Supplementary Figure 2**). Because leiomyosarcoma is characterized by complex karyotypes with extensive CNVs,^2,6,11^ we expected higher InferCNV scores in malignant cells. Clusters 4–6 showed reduced CNV signals that coincided with their lower CytoTRACE scores, indicating that these clusters likely represented benign cell populations (**Figure 1D**).

Cell identities were assigned using canonical, previously reported marker genes.^14^ For example, *CD2* and *CD3D* marked T cells; *VWF* and *ECSCR* marked endothelial cells; and *CD14* and *MRC1*/CD206 marked macrophages (**Figure 1E; Supplementary Table S2**). We also identified differentially expressed genes (DEGs) that defined each specific cluster (**Supplementary Table S3**). Tumor cell clusters were annotated manually based on DEG profiles and pathway enrichment (**Supplementary Table S4**). This revealed substantial heterogeneity and evidence of aberrant differentiation. For instance, one cluster was enriched for neural pathways (cluster 3), whereas others showed enrichment for endocrine progenitor (cluster 5), skeletal muscle (cluster 9), or intestinal smooth muscle associated programs (cluster 11, **Figure 1E, Supplementary Table S4**).

Among non-malignant cells (clusters 4-6), the most common cell types were macrophages, T cells, and endothelial cells (**Supplementary Figure 3A-B**). Most tumors contained macrophages expressing immunosuppressive markers such as *MRC1* and *CD163* (**Supplementary Figure 3C).** In tumors with higher T-cell infiltration, many T cells expressed exhaustion markers including PD1 and CTLA4 (**Supplementary Figure 3D**). Of the tumors with relatively higher proportions of T cells, the T cells expressed markers of exhaustion such as *PD1* and *CTLA4* (**Supplementary Figure 3E)**. Using established annotation guidelines,^15^ we identified T-cell subsets composed of CD4⁺ T cells (10.5%) and CD8⁺ T cells (11.3%). The overall T-cell compartment was heterogeneous but included effector memory T cells (Tem), naïve CD8⁺ T cells, precursor exhausted T cells, and NKT cells (**Supplementary Figure 4**).

Macrophage populations were further analyzed using a previously defined tumor-associated macrophage (TAM) nomenclature.^16^ Based on this classification, TAMs were grouped into interferon-primed (IFN-TAMs), immune-regulatory (Reg-TAMs), inflammatory cytokine–enriched (Inflam-TAMs), lipid-associated (LA-TAMs), pro-angiogenic (Angio-TAMs), resident tissue macrophage–like (RTM-TAMs), and proliferating subsets (Prolif-TAMs; **Supplementary Figure 5**). Using this framework, soft-tissue leiomyosarcomas (extremity and retroperitoneal) displayed prominent IFN-TAM signatures, whereas uterine leiomyosarcomas did not. Reg-TAMs were also enriched in soft-tissue tumors but largely absent in uterine tumors. Small subsets of Prolif-TAMs and Inflam-TAMs appeared only in uterine leiomyosarcoma, and RTM-TAMs were observed exclusively in retroperitoneal tumors. Uterine leiomyosarcomas were predominantly characterized by Angio-TAMs and LA-TAMs.

Next, we compared differences across sites of disease. Surprisingly, overall proportions of major cell clusters did not show dramatic variation between disease sites (**Supplementary Figure 6**), although subtle site-specific differences emerged. Uterine leiomyosarcomas exhibited higher proportions of cluster 1 (chondroblast-like) and cluster 6 (T cells), while extremity tumors showed a higher proportion of endothelial cells (cluster 4) (**Supplementary Figure 6**). Gene set enrichment analysis (GSEA) revealed that retroperitoneal leiomyosarcomas were enriched for interferon gamma signaling, IL6/JAK/STAT signaling, IL2/STAT5 signaling, and complement activity. Extremity leiomyosarcoma was enriched in interferon alpha and gamma pathways; and uterine leiomyosarcoma showed enrichment of E2F and MYC targets (**Supplementary Figure 7**).

Finally, we examined expression of gene signatures corresponding to three transcriptomic subtypes previously identified in bulk RNA-seq studies.^2,4,5,8^ Unexpectedly, the 16 tumors in our cohort did not segregate into these established categories. Instead, individual tumors frequently expressed markers of multiple subtypes, suggesting that at the single-cell level, multiple transcriptional states may coexist within the same tumor (**Supplementary Figure 8**).

### Two major subtypes of leiomyosarcoma identified from snRNA-seq data

One of the most notable features in the snRNA-seq UMAP is that the tumor cells predominantly form two major clusters (**Figure 2A**). We termed one cluster mesenchymal/dedifferentiated (MES), as it demonstrated higher CytoTRACE scores and increased expression of mesenchymal genes, such as *C7* and *DPT* (**Figure 2B-C**). The second cluster was termed smooth muscle cell (SMC), as it appeared more differentiated by CytoTRACE (**Figure 2B**) and showed higher expression of smooth muscle markers (**Figure 2C; Supplementary Figure 9**). Interestingly, most tumors had nearly all malignant cells within one of these two main clusters with two notable exceptions among the 16 tumors (**Figure 2D-F, Supplementary Figure 10**). The two clusters did not segregate by tumor site (retroperitoneal, uterine, or extremity), grade (intermediate vs. high), or stage (**Figure 2F**).

**Figure 2.**
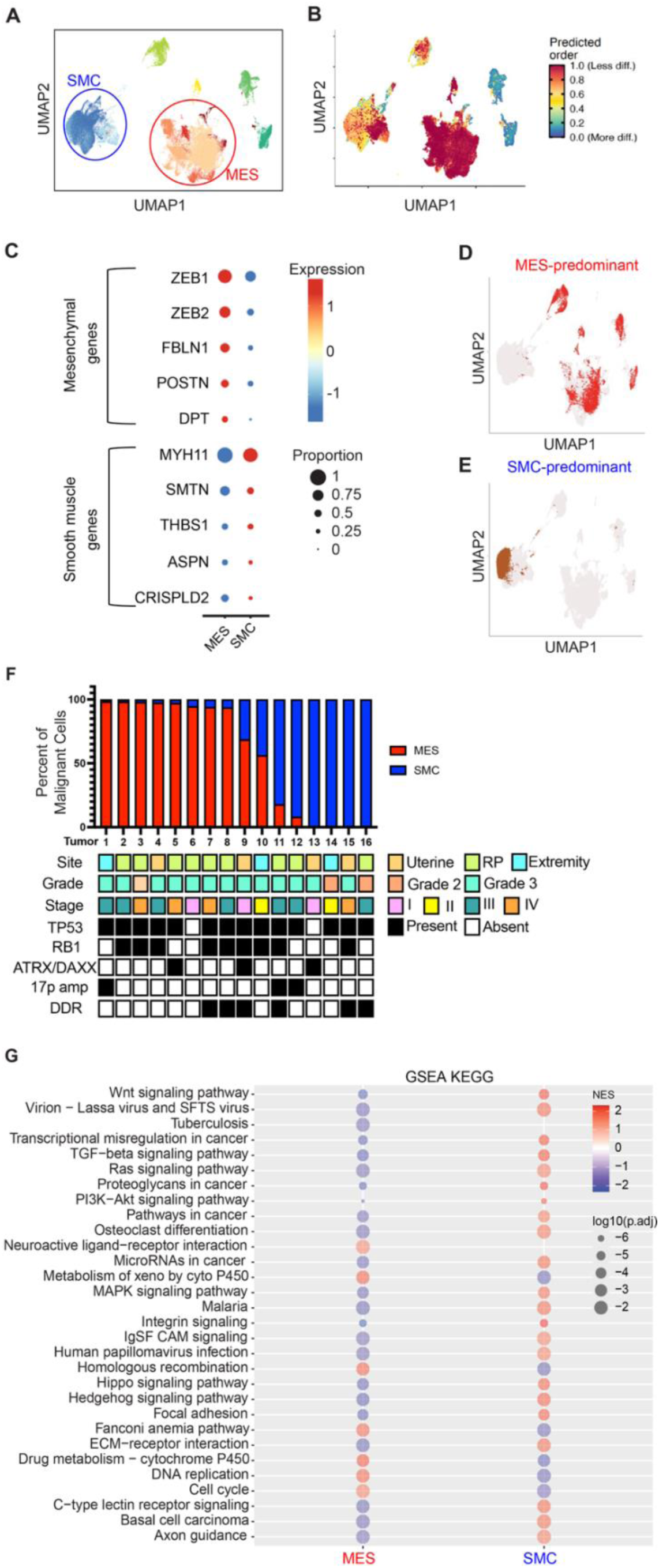
Two novel cellular subtypes of leiomyosarcoma. (A) UMAP of snRNA-seq data demonstrating 2 major tumor cell clusters. MES = mesenchymal, SMC = smooth muscle cell. (B) UMAP with CytoTRACE scores projected onto the snRNA-seq UMAP, demonstrating higher CytoTRACE scores and therefore a more dedifferentiated phenotype in the MES subtype. (C-D) Representative UMAPs of individual tumors with MES- or SMC-predominance. (E) Bubble plot showing enrichment of mesenchymal genes in the MES cluster and smooth muscle genes in the SMC cluster. (F) Quantification of the percent of malignant cells within each tumor that fell within MES (red) or SMC (blue) clusters. Patient, tumor, and genomic characteristics are also indicated for each tumor. Each tumor is represented by its own column. (F) Gene set enrichment analysis comparing MES to SMC.

We also assessed whether genomic profiles could explain these two clusters by performing whole genome sequencing (WGS) (**Supplementary Figure 11; Supplementary Table S5**). WGS identified nearly universal alterations in *TP53* (14/16) and *RB1* (9/16). *ATRX* alterations were detected in 12.5% (2/16), *DAXX* mutation in 6.25% (1/16) and *PTEN* alterations in 18.75% (3/16). In addition, 18.75% (3/16) exhibited 17p amplification encompassing *MYOCD*, which has been implicated in leiomyosarcoma pathogenesis, along with other genes (e.g. *DNAH9, ZNF18, MAP2K4, RNU112P, MIR744, LINC670, ARHGAP*, and *ELOC2*). However, we did not observe any significant differences in genomic profiles between MES- and SMC-predominant tumors. We also did not observe differences in snRNA-seq cluster enrichment based on *ATRX/DAXX* mutation, DNA damage pathway alterations, 17p amplification, or *PTEN* mutation/deletion (**Figure 2F**).

To identify molecular drivers of these two distinct phenotypes identified, we performed GSEA on differentially expressed genes. The MES cluster showed enrichment of neuroactive ligand-receptor interactions, homologous recombination, Fanconi anemia pathway, DNA replication, and cell cycle programs, whereas WNT and TGF-β signaling were specifically enriched in the SMC cluster (**Figure 2G**). Notably, G2M checkpoint and E2F target hallmarks were enriched in the MES cluster, consistent with a more proliferative phenotype.

We also compared metabolic flux using METAFlux^17,18^ to infer metabolic fluxes from our snRNA-seq data. Leiomyosarcoma metabolism is poorly understood, although prior work has shown upregulation of glycolysis, oxidative phosphorylation, glutamate metabolism, and the citric acid cycle.^19,20^ Comparing metabolic flux in MES versus SMC subtypes, we identified enrichment of leukotriene metabolism, cysteine and methionine metabolism, aminoacyl tRNA synthesis, carnitine shuttle, and keratin and chondroitin sulfate degradation in the MES subtype, and enrichment of glycine/serine/threonine metabolism, fatty acid elongation, beta oxidation, and nicotinamide metabolism in the SMC subtype (**Supplementary Figure 12**).

### Immunosuppressive macrophages and exhausted T cells enriched in leiomyosarcomas

We compared the tumor microenvironments of MES- versus SMC-predominant tumors. MES-predominant tumors showed significantly greater immune cell infiltration, particularly macrophages (**Figure 3A-B; Supplementary Figure 13**). The MES cluster also demonstrated higher *IL7* expression compared with SMC (**Supplementary Figure 13**). Macrophages in MES-predominant tumors expressed high levels of immunosuppressive macrophage markers such as *MRC1* and *CD163* (**Figure 3C**). Analysis of targetable immune checkpoints revealed that MES tumor cells expressed higher levels of *CD24* (**Figure 3D, F**) and its receptor *SIGLEC10* (**Figure 3G**), while SMC tumor cells expressed higher levels of *LRP1* (**Figure 3E, H**). The snATAC-seq data revealed open chromatin at the *CD24* locus in MES tumor cells and at the *LRP1* locus in SMC tumor cells, suggesting that expression of these checkpoints may be epigenetically regulated (**Figure 3I)**.

**Figure 3.**
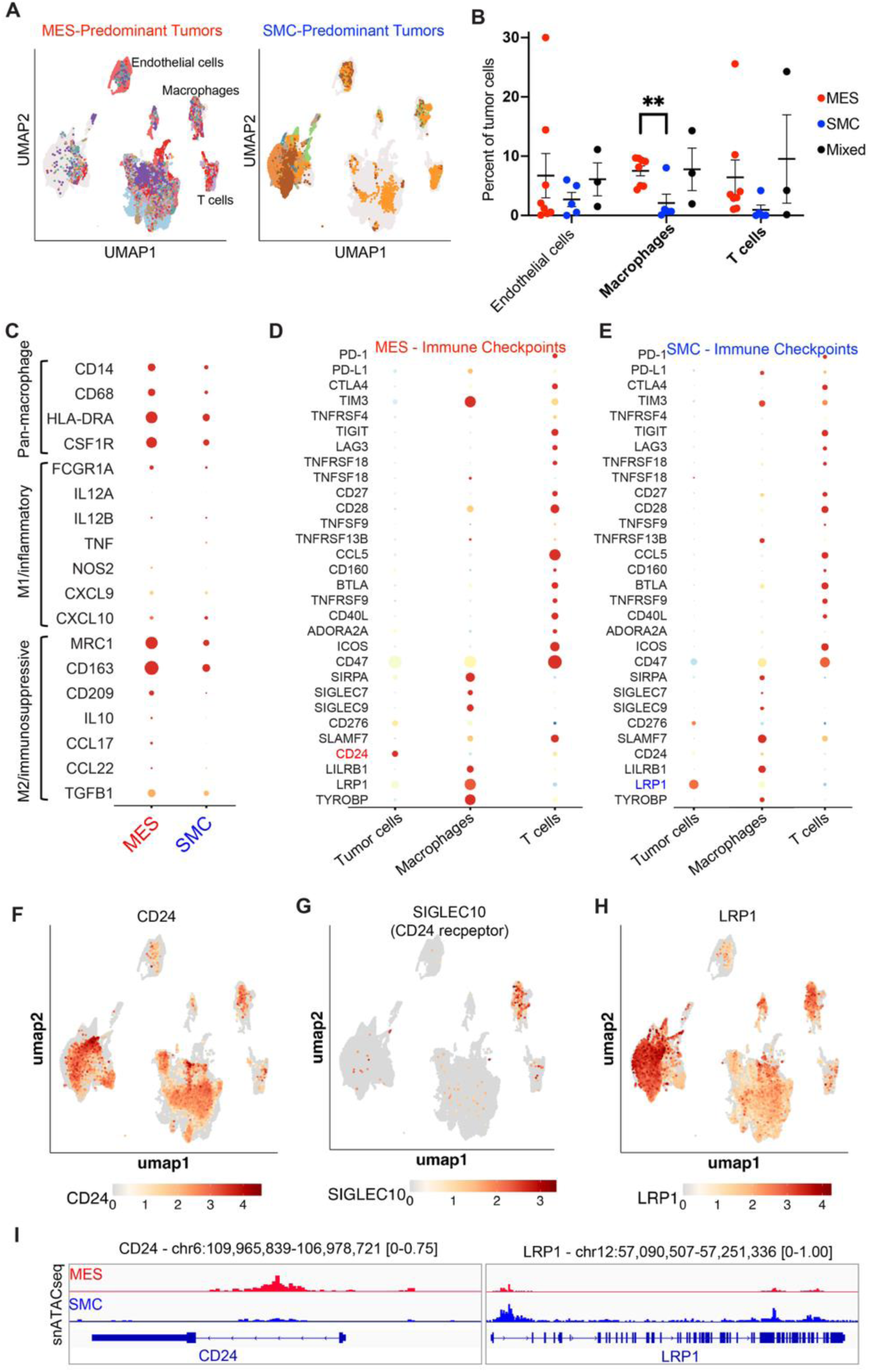
The leiomyosarcoma tumor microenvironment. (A) UMAPs of snRNA-seq data demonstrating differences between MES- versus SMC-predominant tumors. (B) Percent of cells from snRNA-analysis in each of the 3 benign clusters. Each dot represents a single tumor/patient, and bars represent means ± SEM. (C) Bubble plot showing enrichment of macrophage markers in MES vs SMC tumor microenvironments. (D-E) Bubble plots showing expression of T cell and macrophage checkpoints in MES-predominant tumors (D) and SMC-predominant tumors (E). (F-G) UMAP showing expression of *CD24* (G) and *LRP1* (H). (H) Track plots from snATAC-seq data demonstrating open chromatin at *CD24* locus in MES cells and *LRP1* locus in SMC cells.

We applied CellChat to infer interactions between cell populations based on receptor–ligand expression.^21^ The strongest interactions occurred between MES and endothelial cells, MES and fibroblasts, and fibroblasts and endothelial cells (**Supplementary Figure 14**). Prominent autocrine interactions were observed in the MES and fibroblast clusters (**Supplementary Figure 14**). The most upregulated interactions involved collagens, laminin, FN1, ADGRL, tenascins, PTPR, NCAM, FGF, NETRIN, CADM, and PTN. Comparing MES and SMC subtypes, MES was enriched for FN1, ADGRL, tenascin, SEMA3, visfatin, NRXN, EPHA, PTPR, FGF, netrin, CADM, and PTN pathways, whereas SMC was enriched for NCAM1. Overall, MES tumor cells demonstrated higher and more diverse interactions with other cell types than SMC tumor cells.

### Spatial transcriptomics identifies unique spatial niches in MES- and SMC-predominant tumors

Recent technological advances allow spatial characterization of tumor heterogeneity and tumor–microenvironment interactions. Using 10x Xenium *in situ* spatial transcriptomics, we validated MES and SMC predominance in a larger cohort and examined spatial relationships with immune subsets. We designed a custom panel of 480 genes informed by cluster DEGs from the snRNA-seq data, including MES and SMC signature genes and markers of T cells, B cells, NK cells, myeloid cells, fibroblasts, endothelial cells, and neurons (**Supplementary Table S6**). This panel was applied to leiomyosarcoma tissue microarrays (TMAs) comprising 328 tissue cores from 128 patients (**Figure 4A–D; Supplementary Figure 15**). Matched primary and metastatic samples were available for 33 patients. Cells from all TMA slides were combined and clustered, and DEGs were used to annotate clusters (**Figure 4E**). Consistent with snRNA-seq, most tumors contained almost exclusively MES or SMC cells, though many TMA samples demonstrated fibroblast predominance (**Figure 4F; Supplementary Figure 15**), likely reflecting pre-treatment with chemotherapy and/or radiation, which commonly induces fibrosis.^22–24^

**Figure 4.**
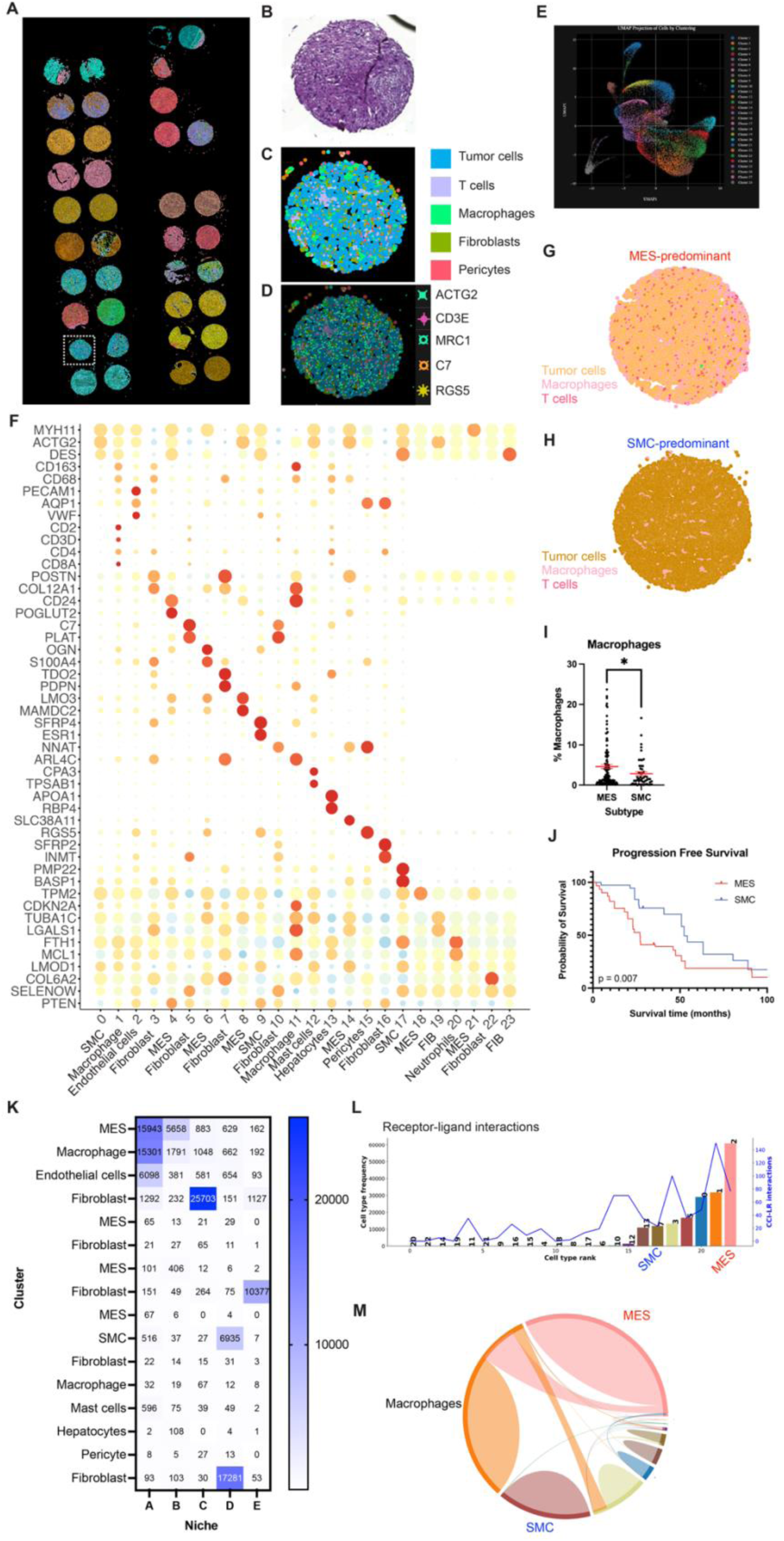
Spatial transcriptomics identifies MES cluster is associated with immunosuppressive microenvironment and worse outcomes. (A-D) Representative image of a leiomyosarcoma TMA analyzed by Xenium. Panels B-D show enlargement of one tissue core’s H&E staining (B), cell annotation (C), and marker gene expression (D). (E) UMAP showing the cell clusters identified. (F) Bubble plot showing maker genes that were used to define each cluster. (G-H) Representative images of MES- and SMC-predominant tissue cores. (I) Quantification of the percent of cells that are macrophages in MES-versus SMC-predominant tumors. *P value < 0.05. (J) Kaplan Meier plot of progression-free survival in metastatic leiomyosarcoma based on MES- versus SMC-predominance. (K) Heatmap of niche analysis demonstrating which clusters were found in each of the 5 niches (A-E) identified. (L) Number of receptor-ligand interactions ranked by cluster. (M) Circos plot demonstrating frequency of receptor-ligand interactions between clusters.

We assessed spatial relationships between these subtypes and infiltrating immune cells. This revealed an enrichment in immunosuppressive macrophages and exhausted T cells in MES tumors compared to SMC tumors (**Figure 4G-I**). MES predominance was associated with significantly worse progression-free survival in patients with advanced/metastatic leiomyosarcoma (**Figure 4J**). In the initial 16-patient single-cell cohort, no differences in outcomes were observed, likely due to small sample size (**Supplementary Figure 16**).

Spatial niche analysis showed that MES tumor cells occupied niches co-populated by macrophages and endothelial cells, whereas SMC tumor cells were associated with fibroblast-enriched niches (**Figure 4K**). Receptor–ligand analysis revealed enriched interactions between MES tumor cells and macrophages (**Figure 4L–M**), with THBS1–CD47 identified as the top receptor–ligand pair (**Supplementary Table S7**).

### snATAC-seq identifies distinct epigenomic profiles in MES and SMC clusters

Given the marked differences in cell phenotypes, we reasoned that underlying epigenomic variation may drive the observed duality. snATAC-seq integration revealed that epigenomic profiles segregated more strongly by snRNA-seq-defined clusters than by tumor of origin (**Figure 5A**) than by patient/tumor sample (**Supplementary Figure 17**), suggesting that these transcriptomic phenotypes reflect underlying epigenetic states. Transcription factor (TF) motif analysis revealed enrichment of NFI family motifs (NFIA, NFIB, NFIC, NFIX) in MES cells and AP-1 family motifs (FOSL2, FOSB, BACH2) in SMC cells (**Figure 5B; Supplementary Table S8; Supplementary Figure 18**). Further, snATAC-seq peaks showed open chromatin at genes that define these different clusters and cell identities (**Figure 5C**). For example, the T cell cluster demonstrated ATAC peaks at *CD3D*. RNA expression similarly showed higher NFI TF expression in MES and higher AP-1 TF expression in SMC (**Figure 5D, Supplementary Figure 19**). Analysis of TCGA data demonstrated that *NFIX* and *FOSL2* are the most highly expressed NFI and AP-1 family members in leiomyosarcoma (**Figure 5E**). These data revealed mutually exclusive tumor groups expressing high *NFIX*/low *FOSL2* or vice versa (**Figure 5F**), supporting a model in which NFI and AP1 family TFs drive distinct, mutually exclusive subtypes in leiomyosarcoma. Further, spatial ATAC-seq, which profiles chromatin accessibility with embedded spatial information,^25^ showed that regions with high NFI activity—particularly NFIB—were spatially associated with *MRC1*-positive macrophage niches in MES-predominant tumors (**Supplementary Figure 20**).

**Figure 5.**
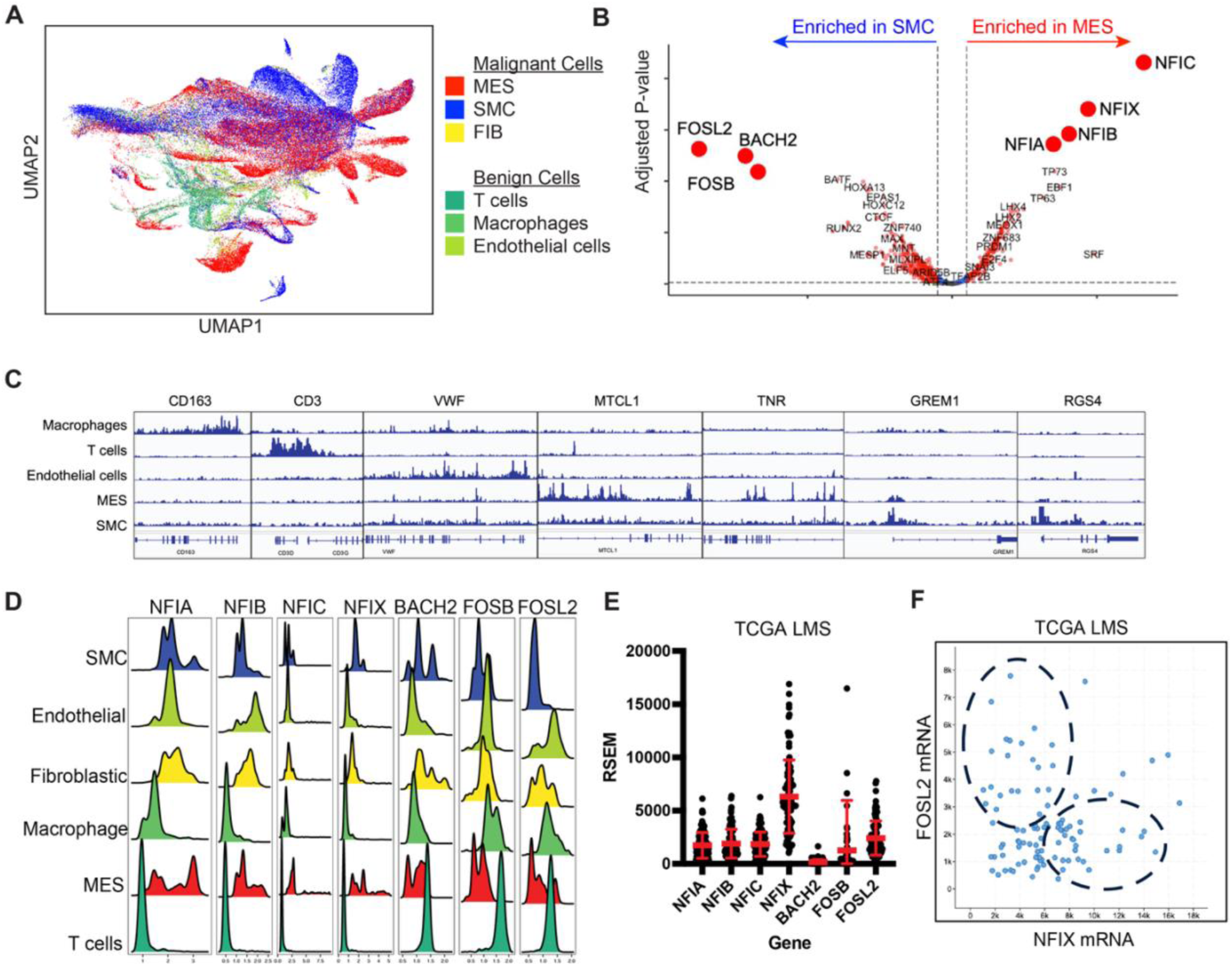
Single cell chromatin openness landscape of leiomyosarcoma demonstrates enrichment of NFI motifs in MES and AP-1 motifs in SMC. (A) UMAP analysis of snATAC-seq with each color representing a different cluster from snRNA-seq analysis. (B) Volcano plot showing enrichment of TF motifs in MES versus SMC. (C) Representative ATAC peaks of genes that define each cluster. (D) Ridge plot showing the distribution of expression of the indicated TFs in the different tumor cell clusters. (E) RNA expression (RSEM) of the indicated TFs in the leiomyosarcoma samples in the TCGA sarcoma dataset. (F) Expression of one of the AP-1 TFs (*FOSL2*) plotted against the expression of one of the NFI TFs (*NFIX*) for the leiomyosarcoma samples in the TCGA sarcoma dataset.

### Leiomyosarcoma cell lines model MES and SMC phenotypes

We utilized human patient-derived leiomyosarcoma cell lines (Leio-012, SK-LMS-1, and SK-UT-1B) and a non-transformed human uterine smooth muscle line (HUtSMC) to functionally validate these findings. Expression of smooth muscle markers, mesenchymal markers, and NFI/AP-1 TFs was assessed by western blotting (**Figure 6A–B**). HUtSMC expressed smooth muscle markers (SMA, MLK1) and minimal mesenchymal markers. Leio-012 expressed SMA and MLK1 at lower levels, expressed AP-1 TFs (FOSL2, BACH2), and showed minimal mesenchymal marker expression, consistent with the SMC phenotype. SK-LMS-01 expressed little or no SMA or mesenchymal markers and low NFI and FOSL2. SK-UT-1B expressed mesenchymal markers (ZEB1, Slug), lacked smooth muscle markers, and expressed high levels of all NFI TFs but minimal AP-1 TFs, consistent with the MES phenotype. Thus, SK-UT-1B best models the MES subtype, whereas Leio-012 approximates the SMC subtype.

**Figure 6.**
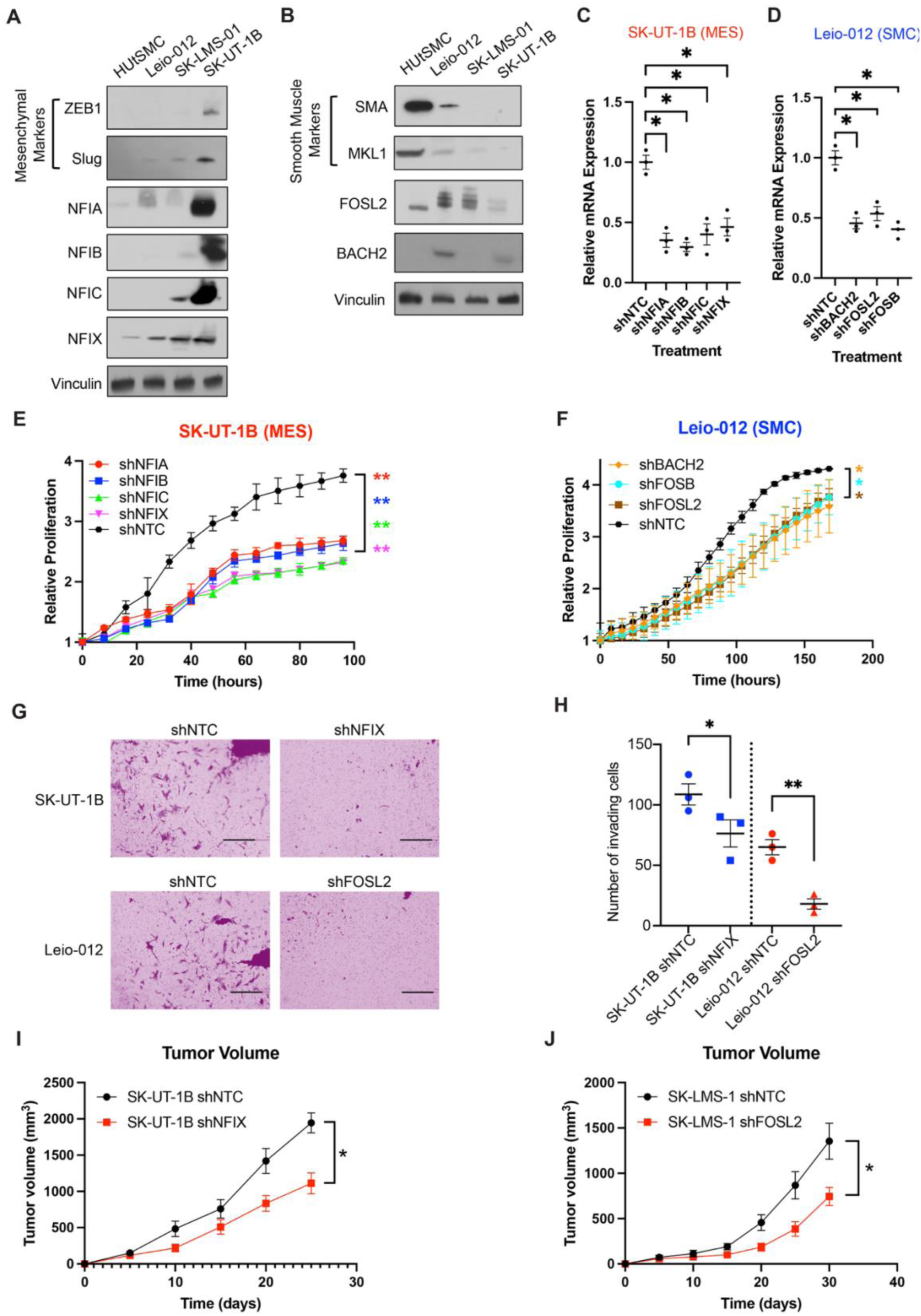
Leiomyosarcoma cell lines recapitulate MES and SMC phenotypes. (A) Western blots showing expression of mesenchymal markers (ZEB1 and Slug) and NFI TFs in untransformed human uterine smooth muscle cells (HUtSMC) versus 3 leiomyosarcoma cell lines (Leio-012, SK-LMS-01, and SK-UT-1B). (B) Western blots showing expression of smooth muscle markers (smooth muscle actin or SMA and MLK1) and AP-1 TFs (FOSL2, BACH2) in the aforementioned cell lines. (C-D) Relative mRNA expression from qRT-PCR showing depletion of the indicated TFs with shRNAs in SK-UT-1B (mimics MES) and Leio-012 (mimics SMC). (E) Relative proliferation of SK-UT-1B cells treated with shRNAs targeting the indicated NFI TFs. (F) Relative proliferation of Leio-012 cells treated with shRNAs targeting the indicated AP-1 TFs. (G) Representative images of invasion assay. (H) Quantification of invasion assay. (I-J) Tumor volume over time following xenografting into immunodeficient nude mice. Timepoints represent means ± SEM for 5 mice. *P value < 0.05; **P value < 0.01.

To test whether these TFs regulate malignant phenotypes, we depleted them using shRNAs (**Figure 6C–D; Supplementary Figure 22; Supplementary Table S9**). NFI depletion reduced proliferation in MES-like SK-UT-1B cells (**Figure 6E**), and AP-1 depletion reduced proliferation in Leio-012 (**Figure 6F**). Depletion of NFIX in SK-UT-1B and FOSL2 in Leio-012 also significantly reduced invasion (**Figure 6G–H**). *In vivo*, NFIX depletion reduced tumor growth in SK-UT-1B xenografts (**Figure 6I**). Leio-012 did not consistently form tumors, so SK-LMS-1 cells were used to test AP-1 function; FOSL2 depletion reduced tumor growth (**Figure 6J**). Together, these results show that NFI TFs drive tumorigenesis and MES phenotypes, while AP-1 TFs drive tumorigenesis and SMC phenotypes.

### Epigenetic driven plasticity between MES and SMC subtypes

We hypothesized that MES and SMC cells might transition between states. Trajectory analysis with Monocle3,^26,27^ which is used to identify potential de-differentiation or evolution trajectories, revealed a potential evolutionary relationship between clusters 7 and 8 (**Figure 7A**), and most tumors contained cells in both clusters (**Figure 7B**), suggesting the potential for plasticity between these clusters. Orthogonal investigation of trajectory using scVelo,^28^ which infers single cell velocity based on the ability to distinguish newly transcribed, unspliced pre-mRNAs from mature, spliced mRNAs by the presence of introns, demonstrated greater velocities within the MES cluster (**Supplementary Figure 23**).

**Figure 7.**
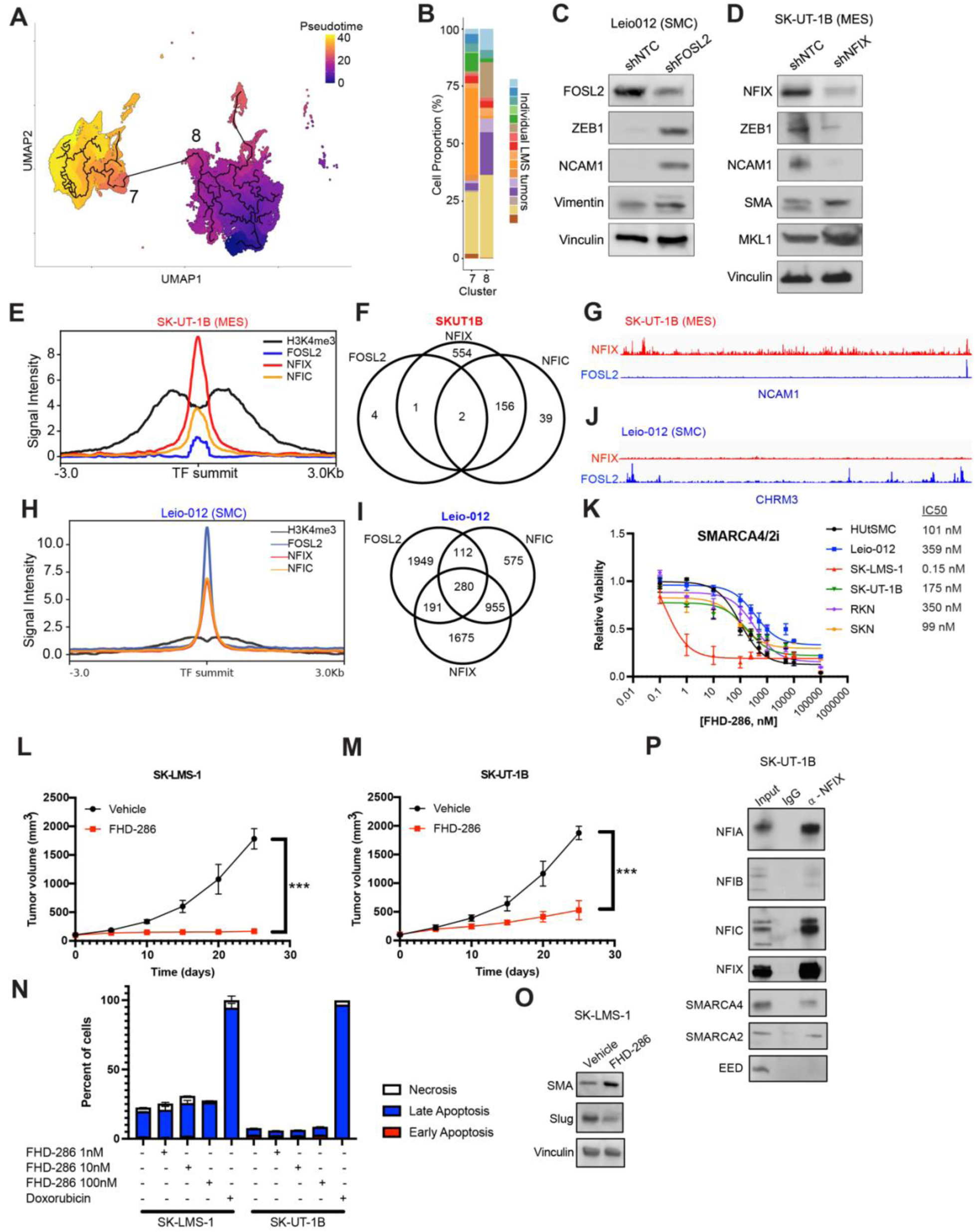
Epigenetic-driven plasticity between MES and SMC subtypes. (A) Pseudotime determined by Monocle 3 is projected onto the snRNA-seq UMAP. A trajectory was noted between clusters 7 (SMC) and 8 (MES). (B) Stacked bar graph demonstrating distribution of cells from individual tumors in different colors in clusters 7 and 8, demonstrating that many of the tumors in the cohort have a population of cells from both clusters 7 and 8. (C) Western blots demonstrating depletion of FOSL2 and subsequent enrichment in mesenchymal markers (ZEB1, NCAM1, vimentin) in an SMC cell line (Leio-012). (D) Western blots demonstrating depletion of NFIX and subsequent decrease of mesenchymal markers (ZEB1, NCAM1) and increase in smooth muscle markers (SMA, MKL1) in a MES cell line (SK-UT-1B). (E) CUT&RUN was performed to assess pathways controlled by each TF. (E) Enrichment of peaks centered on NFIX peaks identified from CUT&RUN data in SK-UT-1B. (F) Venn diagrams showing overlap of peaks identified from CUT&RUN in SK-UT-1B. (G) IGV tracks in a representative MES-defining gene showing the presence of NFIX but absence of FOSL2. (H) Enrichment of peaks centered on FOSL2 peaks identified from CUT&RUN data in Leio-012. (I) Venn diagrams showing overlap of peaks identified from CUT&RUN in Leio-012. (J) IGV tracks in a representative SMC-defining gene showing the presence of FOSL2 but absence of NFIX. (K) Dose response curves with treatment with the SMARCA4/2 inhibitor, FHD-286. IC50 values are shown on the right. (N) Western blots after immunoprecipitating NFIX from SK-UT-1B cells. (L-M) Tumor volume measurements following xenografting of SK-UT-1B (L) and SK-LMS-1 (M) into immunodeficient nude mice and treatment with FHD-286, 1.5 mg/kg given by oral gavage, days 1-5 of 7-day cycles. Treatment started at day 0 on the graph following tumor growth to 100-200 mm^3^. N = 5 mice per group. ***P value < 0.001. (N) Apoptosis assay was performed by staining cells with anti-Annexin V and propidium iodide following treatment with a range of concentrations of FHD-286. Bars represent means ± SEM. (O) Western blotting for smooth muscle (SMA) and mesenchymal (Slug) markers in SK-LMS-1 cells treated with FHD-286. (P) NFIX was immunoprecipitated from SK-UT-1B nuclear extracts and probed for other NFI TFs, SWI/SNF components (SMARCA4/2) and the PRC2 component EED as a negative control.

To test whether NFI and AP-1 TFs drive this plasticity, we depleted FOSL2 in Leio-012 (SMC-like cell line), which increased expression of mesenchymal markers (ZEB1, NCAM1, and vimentin; **Figure 7C**). Conversely, NFIX depletion in SK-UT-1B (MES-like cell line) reduced mesenchymal markers (ZEB1, NCAM1) and increased smooth muscle markers (SMA, MKL1; **Figure 7D**).

To define the target genes and pathways of these TFs, we performed ChIP-seq and CUT&RUN using antibodies targeting each of these TFs and leiomyosarcoma cell lines (**Figure 7E-J**; **Supplementary Table S10**). H3K4me3 served as a positive control with promoter-enriched peaks (**Supplementary Figure 24**). Both NFI and AP-1 TF demonstrated notable enrichment of peaks in intronic and intergenic regions. NFI peaks were enriched for neural-related pathways (e.g., axonogenesis), while AP-1 peaks were enriched for cell differentiation and VEGF signaling (**Supplementary Figure 24**). Taken together, these data demonstrate that MES and SMC cell states are plastic and regulated by distinct TF programs.

### SMARCA4/2 is a potential therapeutic target in leiomyosarcoma

Given the epigenetic basis of these subtypes, we hypothesized that epigenetic therapies might be effective in leiomyosarcoma. A screen of epigenetic drugs (**Supplementary Figure 25**) identified potent low nanomolar sensitivity to the SMARCA4/2 inhibitor FHD-286, particularly in MES-like cell lines (**Figure 7K**). *In vivo*, FHD-286 significantly reduced tumor growth in both SK-LMS-1 (**Figure 7L**) and SK-UT-1B xenografts (**Figure 7M**). FHD-286 did not induce apoptosis (**Figure 7N**) but increased expression of smooth muscle markers (**Figure 7O**), suggesting that it enhances smooth muscle differentiation.

To elucidate the mechanism of sensitivity, we hypothesized that NFI TFs recruit SWI/SNF complexes to remodel nucleosome positioning. NFIX immunoprecipitation pulled down other NFI TFs—consistent with known heterodimerization^29^—as well as SMARCA4 and SMARCA2, but not PRC2 components (**Figure 7P**). This suggests that NFI TFs may drive chromatin remodeling and dedifferentiation by recruiting SWI/SNF complexes.

## DISCUSSION

In summary, this spatially resolved single cell atlas of leiomyosarcoma identifies two novel cellular subtypes of leiomyosarcoma: a mesenchymal/dedifferentiated subtype (MES) and a smooth muscle/differentiated subtype (SMC). These subtypes are driven by specific epigenetic programs with NFI family TFs driving MES subtype and AP-1 family TFs driving the SMC subtype. We further validated functional relevance *in vitro* and *in vivo*, showing that existing leiomyosarcoma cell lines partially recapitulate MES and SMC phenotypes and are sensitive to depletion of NFI and AP-1 TFs. MES predominance is associated with worse outcomes in a cohort of 128 patients. Finally, MES subtypes harbor an immunosuppressive microenvironment with abundance of M2 macrophages.

Our data suggest that NFI and AP-1 family TFs antagonistically regulate phenotypic plasticity in leiomyosarcoma. While the four conserved NFI members (NFIA, NFIB, NFIC, and NFIX) play divergent, context-dependent roles in development and oncogenesis—ranging from glial specification to osteoblast differentiation^30^—their collective role in sarcomagenesis remains largely unexplored. We identified NFI motif upregulation in the mesenchymal (MES) cluster, whereas AP-1 motifs are associated with the PI3K-Akt signaling and smooth muscle contraction pathways enriched in our smooth muscle (SMC) cluster. This highlights a critical role for NFI TFs in the epigenetic reprogramming of leiomyosarcoma. Our data and others^31^ demonstrate that NFI TFs directly interact with the SWI/SNF chromatin remodeling complex, suggesting a potential mechanism by which NFI TFs drive epigenetic reprogramming.

Leiomyosarcoma cells exhibit significant plasticity, likely reflecting the inherent capability of their cell of origin—the smooth muscle cell—to transition between contractile and dedifferentiated fibroblastic states.^32^ This transition, frequently observed in atherosclerosis, is driven by global epigenetic reprogramming.^33^ Notably, our identification of distinct MES and SMC subtypes mirrors the *Zeb2*-mediated phenotypic switching observed in vascular biology, where *Zeb2* loss leads to the enrichment of NFI motifs and a transition to a fibroblastic state.^34^ Similarly, other disease states, such as ischemic cardiomyopathy, are associated with NFI upregulation and downregulation of smooth muscle contraction and PI3K-Akt signaling.^35^ Collectively, these findings suggest that leiomyosarcoma cells are epigenetically poised to navigate these two distinct malignant states, driven by competing transcriptional programs.

Previous bulk transcriptomic studies have consistently identified three transcriptomic subtypes of leiomyosarcoma.^2–8^ We assessed markers of these subtypes and find evidence of expression of the markers throughout many of the tumors analyzed. Thus, the tumors do not fit neatly into one of these subtypes. We hypothesize that this may be due to the presence of multiple cellular subtypes at different proportions in each tumor. Since the clinical presentation and activity of leiomyosarcoma differs by tumor sites,^36,37^ we surveyed and compared different tumors originating from different anatomic sites, though did not find enrichment of any specific anatomical sites in MES or SMC subtypes. This may reflect common pathways and phenotypes among leiomyosarcoma regardless of the site of origin.

Consistent with previous research,^38^ our single-cell atlas confirms that the leiomyosarcoma microenvironment is characteristically immune-poor, dominated by endothelial cells, T cells, and macrophages. We identified a predominance of immunosuppressive M2 macrophages and exhausted T cells (PD-1+/CTLA-4+), providing a biological rationale for the near-zero response rates of traditional immune checkpoint inhibitors (ICI) in leiomyosarcoma.^39,40^ While anti-CD47 therapies have faced clinical hurdles,^41,42^ our data suggest that poor ICI efficacy may stem from targeting suboptimal checkpoints. We identify TIM-3, CCL5, CD24, and LRP1 as promising alternatives; notably, TIM-3 is significantly expressed on leiomyosarcoma macrophages,^43,44^ where it correlates with worse outcomes^45^ and an M2 phenotype via STAT1 signaling.^46^ Given that a rare partial response to anti-TIM-3 therapy was previously observed in a patient with leiomyosarcoma,^47^ our findings support pursuing TIM-3 co-blockade to overcome the suppressive leiomyosarcoma immune landscape.

Limitations of this work include small sample size analyzed by single cell multiome, though we addressed this by expanding the cohort using spatial transcriptomic analysis of leiomyosarcoma TMAs. Furthermore, the samples analyzed by single cell multiome sequencing and spatial profiling represent a small portion of the entire tumor and may not fully reflect intratumoral heterogeneity. Fibroblasts may have been underestimated in single cell multiome sequencing, as we see more fibroblast predominance in the spatial transcriptomic data. This is most likely due to the use of pre-treated samples (e.g. chemotherapy, radiation) in the spatial transcriptomic cohort, as it is widely reported that cancer treatments can cause fibrosis.^22–24^

Overall, we define two clinically relevant, epigenetically regulated leiomyosarcoma cell states with distinct microenvironmental features. These findings provide a framework for subtype-specific therapies and nominate SMARCA4/2 inhibition as a potential strategy for MES-predominant disease. Future work will focus on developing clinically deployable markers and therapeutics targeting these pathways.

## METHODS

### Patients and ethics

This study was approved by the University of Texas MD Anderson Cancer Center (MDACC) Institutional Review Board (protocols PA12-0305, 2022-0278, and LAB04-0890) and was conducted in accordance with the U.S. Common Rule and the Declaration of Helsinki. Clinical and genomic data were obtained following signed informed consent onto prospective institutional protocols or under retrospective review protocols with a limited waiver of authorization, as this is a retrospective project that involves no diagnostic or therapeutic intervention and no direct patient contact. Clinical characteristics including demographic characteristics, disease-associated characteristics (e.g. tumor size, site, etc), and survival were collected from patient electronic health records.

### Patient specimens

Fresh frozen, untreated specimens from 16 patients who underwent surgical resection for leiomyosarcoma were obtained from the MDACC Institutional Tumor Bank (ITB) and utilized for sequencing. The patients had not received chemotherapy or radiation prior to surgery, so the specimens used in this study were not exposed to these treatments. A portion of the same tumor specimen that was used for subsequent single cell sequencing was taken for formalin fixation, paraffin embedding, and hematoxylin and eosin (H&E) staining. All tumor samples were pathologically assessed by an expert pathologist to have a purity of at least 70% and no extensive signs of necrosis.

Tissue microarrays were utilized for spatial transcriptomics and have been previously described.^48–50^ H&E-stained sections were reviewed to confirm the diagnosis, define areas of viable tumor, and select one or more areas (if there was morphological variability) for inclusion in tissue microarrays.

### Nuclei isolation for single-cell sequencing

Nuclei isolation was performed according to the 10x Genomics “Demonstrated Protocol Nuclei Isolation from Embryonic Mouse Brain for Single Cell Multiome ATAC + Gene Expression Sequencing,” CG000366 RevC. Briefly, tissue specimens (approximately 50 mg tissue from each tumor was used) were thawed on ice minced with a scalpel or razor blade on a dish on ice. The minced tissue was added to a 1.5 mL microcentrifuge tube with 500 μL 0.1X lysis buffer, homogenized using a pellet pestle, and incubated on ice for 10 min. Cell/tissue suspensions were mixed 10x with a wide-bore pipette and transferred to a 100 μm Flowmi Cell Strainer and washed with 500 μL wash buffer. Cell suspensions were centrifuged (500g for 5 min at 4 °C) followed by gentle removal of the supernatant without disrupting the cell pellet, another wash with 1 mL wash buffer, centrifuged, and resuspended in wash buffer, and passed through a 40 μm Flowmi Cell Strainer. The concentration and quality of nuclei were assessed using 0.4% trypan blue staining and verification on a hemocytometer. The nuclei were centrifuged again and resuspended in an appropriate volume of chilled 1X Nuclei Buffer (10x Genomics, 2000153) to a concentration of 8000 nuclei/μL. Nuclei were immediately used to prepare snATAC-seq and snRNA-seq libraries.

### Library preparation for 10x Genomics single cell multiome ATAC and gene expression

Single cell multiome sequencing was performed using 10x Genomics Chromium Next GEM Single Cell Multiome ATAC + Gene Expression Reagent Bundle (10x Genomics, Pleasanton, CA, USA) following the manufacturer’s protocol “CG000338 Rev E Chromium Next GEM Single Cell Multiome ATAC + Gene Expression.” The main steps of this protocol include: (1) nuclei transposition, (2) GEM generation and barcoding, (3) post GEM incubation cleanup, (4) pre-amplification PCR, (5) ATAC library construction, (6) cDNA amplification, and (7) gene expression library construction. Briefly, nuclei suspension from previous step with targeted recovery of 10,000 nuclei per sample were mixed and incubated (37 °C for 60 min) with transposition mix. Next, the transposed nuclei were mixed with master mix containing barcoding reagents and loaded onto a Chromium Next GEM Chip J along with Single Cell Multiome GEX Gel Beads and partitioning oil. Gel beads-in-emulsion (GEMs) were generated by delivering nuclei at a limiting dilution to the Gel Beads using the 10x Chromium Controller. The GEMs were captured and cleaned up. Next, the GEMs were dissolved, releasing oligonucleotides containing an Illumina P5 sequence, a 16 nucleotide 10x barcode for ATAC, and a space sequence. Primers containing an Illumina TruSeq Read 1 primer, 16 nucleotide 10x barcode for GEX, and a 12-nucleotide unique molecular identifier, and a 30 nucleotide poly(dT) sequence were also released during this step. These oligonucleotides are mixed with the nuclei lysate containing transposed DNA fragments, mRNA, and master mix containing reverse transcriptase, generating the 10x barcoded transposed DNA for ATAC-seq and 10x barcoded cDNA for RNA-seq. The reaction was quenched with a quenching reagent. GEMs were then broken, and pooled fractions recovered using Silane magnetic beads to purify barcoded products. Barcoded transposed DNA and cDNA were amplified to fill in gaps for subsequent ATAC library construction and cDNA amplification. Next, the snATAC-seq library was constructed by adding P7 and a sample index and performing PCR. Attention was turned to creating the snRNA-seq library, first by amplifying cDNA by PCR to generate sufficient material for library construction, then fragmented and size selected using SPRIselect (Beckman Coulter) to optimize the cDNA amplicon size. P5, P7, i7 and i5 sample indexes, and TruSeq Read 2 primer sequences are added via end repair, A-tailing, adaptor ligation, and PCR, followed by purification with SPRIselect and elution of final libraries.

Next, the libraries were checked for the fragment size distribution using Agilent 4200 Tape Station HS D1000 Assay (Agilent Technologies) and quantified with Qubit Fluorometric dsDNA Quantification kit (Thermo Fisher Scientific). The libraries were sequenced at the MDACC Advanced Technology Genomics Core facility, with each library on a separate lane of HiSeq4000 flow cell (Illumina), with the sequencing targeting above the minimum of 25,000 read pairs per nucleus sequencing depth, format of 100nt and parameters (Read 1—50 cycles, Read 2—50 cycles; with exception while sequencing together with snRNA-seq libraries, Read 1—100 cycles, Read 2—100 cycles).

### Single cell multiome data analysis

Raw multiomic sequencing data were aligned to the human reference genome (Hg38) using Cellranger-Arc v2.0.0, with subsequent annotation of cellular barcodes. Ambient RNA contamination in the snRNA-seq data was mitigated using DecontX,^51^ and after filtering for mitochondrial (removed cells with > 30% mitochondrial RNA) and ribosomal (removed cells with > 10% ribosomal RNA) gene content, doublets were removed with DoubletFinder.^52^ Normalization of snRNA-seq data was carried out using SCTransform, and batch effects were assessed via k-BET and corrected using Harmony and Seurat v5’s integration layers framework, with each layer corresponding to an individual sample. Log-transformed unique molecular identifier counts were used for principal component analysis. Highly variable genes were identified using Seurat, enabling unsupervised clustering, and UMAP was employed for dimensionality reduction and visualization. Differential gene expression was determined using the non-parametric Wilcoxon rank-sum test. Pathway enrichment was evaluated through three distinct methods using differentially expressed genes: Gene Set Enrichment Analysis (GSEA), KEGG pathway analysis, and Reactome pathway analysis. Lineage-specific gene markers were used to conduct cell cluster analysis. Malignant cells were distinguished from non-malignant and immune cells by detecting increased copy number variations (CNVs) through InferCNV.^53^ Additionally, CytoTRACE^13^ was used to infer differentiation states. Monocle3^27^ and scVelo^28^ were employed for trajectory inference. Cluster annotation was performed manually based on expression of certain gene markers (**Supplementary Table 2**).

ArchR was used to process snATAC-seq data.^54^ After removing low quality cells by examining the transcription start site enrichment score and unique nuclear fragments, we used cellular barcode left from snRNA analysis to filter out redundant cells. We used the ArchR gene activity score to calculate gene-level changes in chromatin state for inferential analysis, then integrated with snATAC with matched data. We called differential accessible peaks across groups by testing for statistical differences in the read counts between groups for each peak, using a generalized linear model with binomial link.. Furthermore, ChromVAR^55^ was used to identify enriched TFs on a per-cell basis from sparse chromatin accessibility data.

### Prediction of ligand–receptor interactions

To predict potential ligand-receptor interactions between tumor microenvironment populations and distinct malignant cell states in snRNA-seq data, we employed the R implementation of CellChat v2.1.0^21^ available on GitHub (https://github.com/sqjin/CellChat). A ligand-receptor interaction matrix was generated using a Seurat object as input, with a minimum threshold of 20 cells per group required to infer cell-cell communication. The inferred cellular communication network was extracted as a data frame, allowing for the analysis of intercellular communication at the signaling pathway level, which was further explored through the interactive CellChat explorer.

### Prediction of metabolic flux

METAFlux^17,18^ was used to calculate metabolic fluxes from the snRNA-seq data. METAFlux used the velocity of metabolic reactions and fluxes to predict metabolic reaction activity in our gene expression data.

### Whole genome sequencing

Genomic DNA was extracted from fresh frozen leiomyosarcoma samples using the Qiagen DNeasy Blood & Tissue Kit and sent to GENEWIZ for whole genome sequencing with an average sequencing depth of 20x.

### Spatial transcriptomics

The 10x Genomics Xenium *in situ* platform was utilized for spatial transcriptomics. We designed a custom 480-gene panel using DEGs from the snRNA-seq data and well-established markers of different immune cell subsets. We also included genes that define different leiomyosarcoma subsets from bulk transcriptomic data from prior studies.^2,4,5^ A list of genes is shown in **Supplementary Table S6**. We utilized previously reported leiomyosarcoma TMAs, which are clinically annotated with data including, but not limited to, grade, stage, progression-free survival, overall survival, and response to chemotherapy,^48,49^ and 5 μm formalin-fixed, paraffin-embedded sections were placed onto a Xenium slide. Tissues underwent deparaffinization and permeabilization, then the custom probe panel at 10 nM concentration was hybridized overnight at 50° C. Slides were washed, then probes were ligated to seal the junction between probe regions that had hybridized to RNA at 37° C for 2 hours, followed by amplification of ligation products at 37° C for 2 hours. Tissues were washed, chemically quenched of background fluorescence, and nuclei stained with DAPI. Slides were loaded onto a Xenium Analyzer. The Xenium cell segmentation algorithm used DAPI images that were acquired and calculated in a 15-μm radius from the nucleus outward or until another cell boundary was reached. The on-instrument pipeline output included files containing the feature-cell matrix, the transcripts, and a CSV file of the cell boundaries (a differentially enriched gene list for each cluster). Further downstream analysis and data integration for all samples were performed off-instrument and visualized using 10x Genomics Xenium Explorer (version 3). After completing the run, H&E staining was performed on the slide. A Keyence microscope was used to obtain images of the stained slides.

Spatial niche analysis was performed on the Xenium spatial transcriptomics dataset using Seurat. Cell identities were first defined based on unsupervised clustering, with cluster assignments stored in the Seurat clusters metadata field. All cells present in the selected field-of-view were included in downstream niche analysis. A niche assay was constructed using the BuildNicheAssay function. For each cell, a local neighborhood was defined by identifying its 30 nearest spatial neighbors (neighbors.k = 30). Neighborhoods were characterized by the composition of Seurat clusters within each local neighborhood and subsequently grouped into 15 distinct spatial niches (niches.k = 15) based on similarity in cluster composition. Spatial distributions of both cell types and inferred niches were visualized using ImageDimPlot, to projects cells on to their original tissue coordinates. Cells were colored either by their Seurat cluster identity or by their assigned niche identity. Custom color palettes were applied to enhance visual distinction between niches. Further, to ensure the data integrity and absence of any missing values, a contingency table was generated between cell-type clusters and spatial niches to summarize the number of cells from each Seurat cluster within each niche. This table was converted to a matrix format and exported as a CSV file for downstream statistical analysis. Finally, the processed Seurat object containing niche assignments with metadata formation was saved as an RDS file for downstream analyses.

### Spatial ATAC-seq

Fresh frozen leiomyosarcoma tissue (same as used for single cell multiome sequencing) was embedded in optimal cutting temperature (OCT) compound and cryosectioned into 10um slides, affixed to slides, and stored at -80° C. Samples were shipped to AtlasXomics (New Haven, CT) in dry ice for spatial-ATAC-seq. The frozen slides were warmed at room temperature for 10 min. The tissue was then fixed with formaldehyde (0.2% for 5 min) and quenched with glycine (1.25 M for 5 min) at room temperature. After fixation, the tissue was washed twice with 1 mL of 1x PBS, cleaned in deionized water, and dried. The samples were then used for spatial ATAC-seq after optimization of assay conditions and flow testing. Deterministic Barcoding in Tissue sequencing (DBiT-seq), a commercial, first-generation, 50-channel platform (Portal50, AtlasXomics), was used for spatial ATAC-seq.

### Cell culture

HUtSMC, SK-LMS-01, SK-UT-1B, BJ, and HDFn were obtained from ATCC. HEK293T was obtained from the MD Anderson cell bank. Leio-012 cell were derived as described.^56^ RKN and SKN were obtained from JCRB Cell Bank. All cell lines were authenticated using STR analysis at the MDACC Cytogenetics and Cell Authentication Core. Cell lines were routinely tested for Mycoplasma using the Mycoplasma PCR Detection kit (abm). All cell lines were grown in DMEM (HyClone, SH30243.01) with 10% fetal bovine serum (Gibco) and 1% penicillin/streptomycin (HyClone, SV30010).

For proliferation and growth curves, an Incucyte was used (Sartorius). For invasion assays, 5x10^4^ cells were seeded in serum-free media onto BioCoat Matrigel Invasion Chambers (Corning, 8 μm pore). Complete media (10% FBS) was added into well below the chamber. After 24-hour incubation, a cotton swab was used to remove cells on top of the membrane, and the cells on the lower surface of the membrane were stained with crystal violet and quantified by microscopy with manual counting.

Apoptosis assay was performed by harvesting cells, washing once with PBS, resuspending in Annexin V binding buffer (10 mM HEPES pH 7.4, 140 mM NaCl, 2.5 mM CaCl_2_), and staining cells with anti-Annexin V and propidium iodide (Biolegend). Data were acquired on a Gallios flow cytometer (Beckman Coulter).

Lentiviruses were generated by transfecting a T25 of ∼70% confluent HEK-293T cells with 2 μg pLKO.1 vector, 1 µg pVSV-G, 1 µg psPAX2, and Lipofectamine 3000 (Thermo Fisher). Fresh medium was applied at 24 hours post-transfection, and virus was harvested 24 hours later, clarified by centrifugation at 1000g x 5 minutes and filtration through a 0.45 μm filter to remove cell debris, and diluted 1:4 with complete medium containing 10 μg/mL polybrene. Target cells were transduced at ∼50% confluence for 24 hours, then given fresh media for 24 hours, then 1 µg/mL puromycin was added for 3-5 days.

Chemicals used in this study include: puromycin (InvivoGen), polybrene (InvivoGen), FHD-286 (MedChemExpress), panobinostat (MedChemExpress), vorinostat (MedChemExpress), tazemetostat (MedChemExpress), azacitidine (MedChemExpress), decitabine (MedChemExpress), A-845 (MedChemExpress), JQ1 (MedChemExpress), ZEN-3694 (MedChemExpress), bomedemstat (MedChemExpress), SNDX-5613 (MedChemExpress), MI-503 (MedChemExpress), gintemetostat (MedChemExpress), GSK-J4 (MedChemExpress).

### Molecular Biology

Human shRNAs (Horizon GIPZ Lentiviral shRNA) and cDNAs (Horizon Precision LentiORF collection) for the following genes were obtained from the MDACC Functional Genomics Core: NFIA (clone ID V3LHS_307343, mature antisense TCTTCTTTTGACATACGCT), NFIB (clone ID V3LHS_366521, mature antisense AAAGTACTTGCGTTTTCGA), NFIC (clone ID V3LHS_329715, mature antisense TGTCCATCTCTGTCTTCTT), NFIX (clone ID V3LHS_343918, mature antisense TTGATTGTGACTCCAATGT), BACH2 clone (ID V3LHS_640501, mature antisense AGCTTGTGCATTTTAATCA), FOSB (clone ID V2LHS_85028, mature antisense TCTTCCTCCAACTGATCTG), FOSL2 (clone ID V3LHS_637338, mature antisense CAAGGAGTCTGATGATTGG), and non-targeting control (mature antisense ATCTCGCTTGGGCGAGAGTAAG). Plasmids were used to generate lentivirus, as described above.

The CRISPR/Cas9 system was used to knockout TFs in leiomyosarcoma cell lines. Guides were designed using CRISPOR,^57^ custom oligos were purchased from Sigma and cloned into lentiCRISPRv2 (a gift from Feng Zhang, Addgene plasmid #52961). Plasmids were used to generate lentivirus, which was used to transduce target cells, as described above. Transduced cells were selected with puromycin (1 μg/mL) for 72 hours and then allowed to recover for 24 hours in fresh medium.

### Western blotting

Cells were lysed in RIPA buffer (Boston Bioproducts, BP-115) with protease and phosphatase inhibitors (Halt, Thermo, 1861282). Proteins were separated by SDS-PAGE (Biorad pre-cast gels), transferred to PVDF membrane (Biorad Trans-Bot turbo 0.2μm PVDF) using Biorad Trans-Blot Turbo Transfer System, blocked with Biorad EveryBlot Blocking Buffer for 5-10 minutes, and probed overnight in 4°C with primary antibodies diluted in EveryBlot Blocking Buffer. Horseradish peroxidase-conjugated rabbit and mouse secondary antibodies (CST 7074 and 7076) were applied for 2-3 hours at room temperature. Primary antibodies used include: NFIA (CST, 69375, 1:1000), NFIB (CST, 24903, 1:1000), NFIC (Abcam, ab245597, 1:1000), NFIX (Thermo, MA5-946983, 1:1000), BACH2 (CST, 80775, 1:1000), FOSB (CST, 2251, 1:1000), FOSL2 (CST, 19967, 1:1000), smooth muscle actin (CST, 19245, 1:1000), MKL1, (Proteintech, 21166-1-AP, 1:1000), ZEB1 (CST, 3396, 1:1000), Slug (CST, 9585, 1:1000), NCAM1 (CST, 99746, 1:1000), lamin B1 (Abcam, ab229025, 1:1000), alpha tubulin (Santa Cruz sc-32293, 1:5000), beta actin (Sigma, A2228, 1:5000), and vinculin (Santa Cruz, sc-59803, 1:1000).

### Immunoprecipitation

For NFIX immunoprecipitation, cells were lysed in hypotonic buffer (20 mM HEPES pH 7.4, 10mM KCl, 2 mM MgCL2, 1mM EDTA, 1mM EGTA, 0.1% IGEPAL, 1mM DTT, and 1X Halt protease and phosphatase inhibitor cocktail (Thermo, 78440)). Nuclei were pelleted by centrifuging at 720g for 5 min. Nuclei pellets were washed with the same hypotonic lysis buffer and lysed in immunoprecipitation buffer (50 mM Tris-HCl pH 7.5, 150 mM NaCl, 1mM EDTA, 1% Triton X-100, 1 mM DTT, and 1X Halt protease and phosphatase inhibitor cocktail). Supernatants (nuclear extracts) were collected by centrifugation at 17,000g for 15 min and incubated overnight with anti-NFIX antibody (Sigma-Aldrich, SAB1401263, 1:100). Protein G Dynabeads (Invitrogen, 10004D) were added and incubated for 4 hours at 4 °C. The precipitates were washed three times with immunoprecipitation buffer, boiled in sample buffer, and utilized for western blotting.

### Quantitative reverse transcription polymerase chain reaction (qRT-PCR)

For gene expression analysis by qRT-PCR, RNA was isolated from cells using Qiagen RNeasy kit. Reverse transcription was performed using SuperScript VILO (Thermo Fisher). Samples were amplified on a QuantStudio 6 (Thermo Fisher). The average C_T_ value of three housekeeping genes (*RRN18S, GAPDH,* and *ACTB*) was subtracted from each experimental C_T_, then 2−ΔCT values were normalized to controls (non-targeting shRNA control) and compared. Additionally, the ΔΔC_T_ method was employed to calculate the fold change in gene expression. Primer sequences are provided in **Supplementary Table 8**.

### Immunofluorescence

Cells were plated onto glass coverslips in 12-well plates and fixed with 4% paraformaldehyde in PBS. Fixed cells were then blocked for 30 min in 3% bovine serum albumin (BSA) and 0.1% triton X-100 in PBS (PBSTx + BSA). Primary antibodies were incubated in PBSTx + BSA for 1 h at room temperature and washed three times in PBSTx, followed by secondary antibody incubation in PBSTx + BSA for 30 min at room temperature and two washes with PBSTx. Cells were counterstained with DAPI and mounted on glass slides with Prolong Gold antifade medium (Invitrogen). Image acquisition was performed using a Keyence BZ-X microscope or a Confocal Zeiss LSM880 confocal microscope with Airyscan super-resolution.

Antibodies used for immunofluorescence include: NFIA (CST, 69375), NFIB (CST, 24903), NFIC (Abcam, ab245597), NFIX (Thermo, MA5-946983), and BACH2 (CST, 80775), FOSL2 (CST, 19967),; all were used at 1:200 dilution in PBSTx + BSA. Alexa fluor-conjugated secondary antibodies were used (Invitrogen, 1:350).

### Xenografting

SK-LMS-1 and SK-UT-1B cells (5 x 10^6^) were injected into the flanks of nude mice (NU/J, strain #002019, Jackson Laboratory). Calipers were used to measure tumor size, and tumor volume was calculated according to the formula (a × b^2^)/2, where “a” was the longest diameter and “b” was the shortest diameter of the tumor.

For treatment with FHD-286, mice were xenografted with SK-LMS-1 and SK-UT-1B cells as above, and treatment with 1.5 mg/kg FHD-286 (by oral gavage, dissolved in corn oil, days 1-5 every 7-day cycle) was started when tumors reached approximately 100 mm^3^.

### Chromatin immunoprecipitation (ChIP)

ChIP was performed as previously described previously.^58^ Cells (5 million per reaction) were cross-linked using 1% paraformaldehyde for 10 minutes at 37°C. Reactions were quenched by 0.125 M glycine for 5 minutes. Cells were then washed with PBS and stored at –80°C. Cells were thawed on ice the next day and lysed with ChIP harvest buffer with protease inhibitors (12 mM Tris-Cl, 0.1x phosphate buffered saline (PBS), 6 mM EDTA, 0.5% sodium dodecyl sulfate (SDS) for 30 minutes on ice. Sonication was performed using the Branson Sonifier 250 to achieve a DNA shear length of 200 to 500 bp. Extracts were then incubated overnight with their respective antibody-protein G Dynabead mixture (Invitrogen). Antibodies used for ChIP include: NFIA (CST, 69375), NFIB (CST, 24903), NFIC (Abcam, ab245597), NFIX (Thermo, MA5-946983), BACH2 (CST, 80775), FOSL2 (CST, 19967). Immune complexes were then washed 5 times with RIPA buffer, twice with RIPA-500 (RIPA with 500 mM NaCl), and twice with LiCl wash buffer (10 mM Tris-HCl, pH 8.0, 1 mM EDTA, pH 8.0, 250 mM LiCl, 0.5% NP-40, 0.5% DOC). Elution and reverse crosslinking were performed in direct elution buffer (10 mM Tris-Cl, pH 8.0, 5 mM EDTA, 300 mM NaCl, 0.5% SDS), with addition of 1 µL of RNase (20mg/mL), and 5 µL Proteinase K (20mg/mL), and incubating reactions at 65°C for 12 hours. DNA was cleaned up using SPRIselect beads (Beckman-Coulter).

Libraries were prepared using NEBNext Multiplex Oligos for Illumina (E6611A). Libraries were pooled and sequenced on Illumina NextSeq 550 at the MDACC Advanced Technology Genomics Core.

### CUT&RUN

Cleavage under targets and release using nuclease (CUT&RUN) was performed following EpiCypher’s CUTANA protocol and reagents. Briefly, cells were harvested with 0.05% trypsin, and 1x10^6^ cells were used per reaction. Cells were bound to concanavalin-conjugated paramagnetic beads. Antibodies were added and allowed to bind overnight. Antibodies used included: NFIA (CST, 69375), NFIB (Bethyl, A303-566A-T), NFIC (Abcam, ab245597), NFIX (Thermo, MA5-946983), BACH2 (CST, 80775), FOSL2 (CST, 19967), rabbit IgG (Abcam ab171870), H3K4me3 (Abcam, ab8580). Stop buffer was added with E. coli DNA spike-in. pAG-MNase was added and activated with CaCl_2_. DNA was purified using SPRIselect, and concentration checked by Qubit fluorometer per manufacturer’s instructions. Library prep was performed using 5 ng input DNA and CUTANA CUT&RUN Library Prep Kit. Libraries were pooled (each sample diluted to 2nM) and sequenced on Illumina NextSeq 550 at the MDACC Advanced Technology Genomics Core.

### Statistical analysis of *in vitro* experiments

Statistical analyses were performed using GraphPad Prism (version 9.5.0 or higher, RRID:SCR_002798) and R (version 4.2.2 or higher). Survival was assessed using the Kaplan Meier method. Cox proportional hazards modeling was performed using survival package in R.

For *in vitro* experiments, the data from 3 biological replicates are presented as mean ± SD, and SD represents the deviation between the biological replicates. Graph Pad Prism (version 10.6.1) and R were used for statistical analyses.

Graphics for schematics in the figures were created using Biorender.com.

## Acknowledgements

The authors thank Dona and Gene Scott, the QuadW Foundation, and the National Leiomyosarcoma Foundation (NLMSF) for supporting this project. The authors also acknowledge support from the MD Anderson Cancer Center Support Grant (P30 CA016672). This research is supported by The MD Anderson Cancer Center Sarcoma SPORE Grant P50CA302482. Research reported in this publication was supported by the National Cancer Institute of the National Institutes of Health under Award Number P50CA272170. The content is solely the responsibility of the authors and does not necessarily represent the official views of the National Institutes of Health. RAD is supported by NIH grant T32 CA009666 and a Conquer Cancer - Endowed Young Investigator Award in Honor of Grant R. and Victoria A. Merryman. Any opinions, findings, and conclusions expressed in this material are those of the author(s) and do not necessarily reflect those of the American Society of Clinical Oncology® or Conquer Cancer®. ENH is supported by the Paul Calabresi Career Development Award for Clinical Oncology as funded by the NCI/NIH through grant 5K12CA088084, as well as the BMS foundation through Robert A Winn Career Development Award. The results shown here are in part based on data that were generated by the TCGA Research Network: http://cancergenome.nih.gov/. We acknowledge the TCGA Research Network, including the specimen donors and research groups, for their contributions.

## Abbreviations

ChIP: chromatin immunoprecipitation
CNV: copy number variation
CUT&RUN: cleavage under targets & release using nuclease
DEG: differentially expressed gene
GEM: gel beads-in-emulsion
GSEA: gene set enrichment analysis
H&E: hematoxylin and eosin
MDACC: MD Anderson Cancer Center
MES: mesenchymal subtype
qRT-PCR: quantitative reverse transcription polymerase chain reaction
SMC: smooth muscle cell subtype
snATAC-seq: single nucleus assay for transposase-accessible chromatin with sequencing
snRNA-seq: single nucleus RNA sequencing
TF: transcription factor
TMA: tissue microarray

