## Supplemental Figures for "Spatially-Resolved Multiomic Atlas of Leiomyosarcoma Identifies Two Clinically Relevant Epigenetically-Driven Cell States"

SUPPLEMENTARY DATA

**Table S1. Number of cells per sample.**

**Table S2. Genes used for cluster annotations.**

**Table S3. Differentially expressed genes for each cluster.**

**Table S4. Cluster annotations.**

**Table S5. Whole genome sequencing data.**

**Table S6. List of genes included in Xenium spatial transcriptomics custom panel.**

**Table S7. Top ligand-receptor interactions.**

**Table S8. TF motif enrichment in MES versus SMC.**

**Table S9. Primer sequences for qRT-PCR.**

**Table S10. ChIP-seq and CUT&RUN experimental parameters.**

**Supplementary Figure 1. Histologic validation of leiomyosarcomas.**
Micrographs of H&E-stained slides for each of the 16 leiomyosarcoma samples.

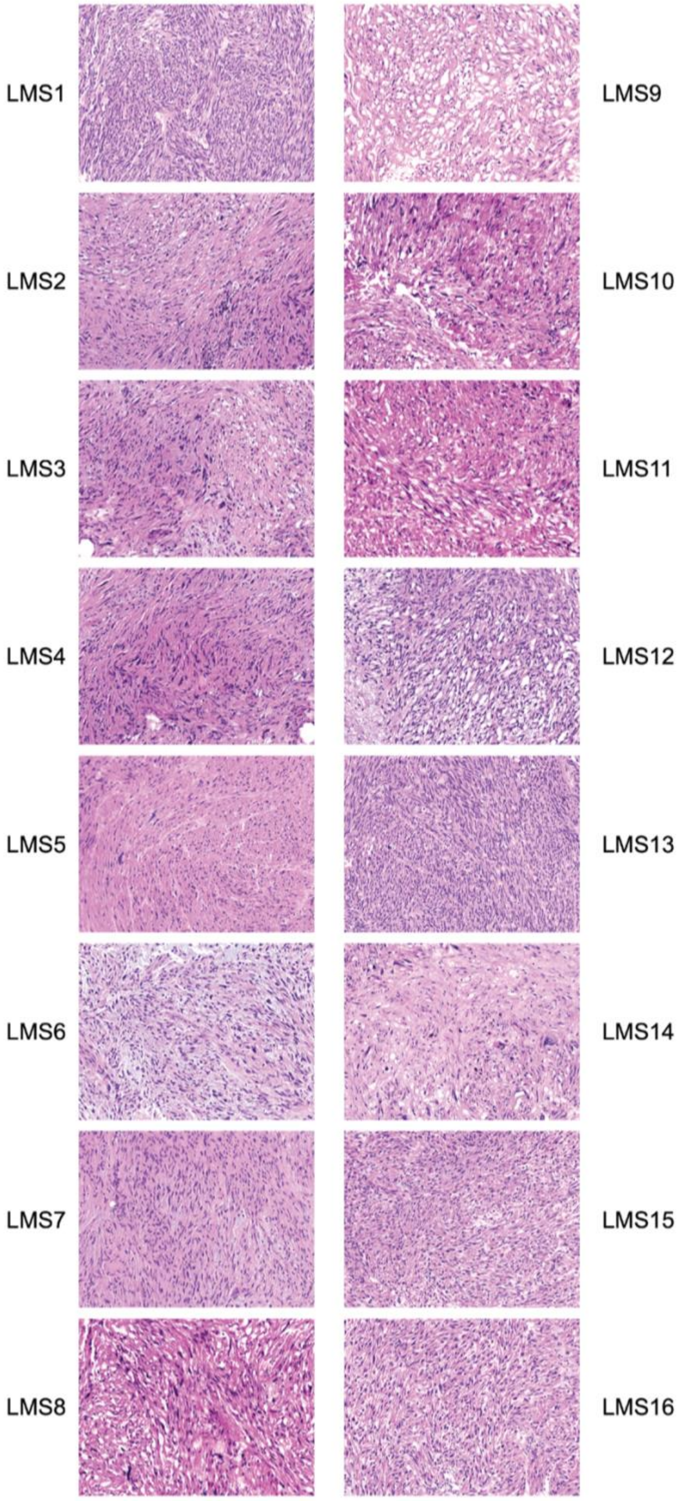

**Supplementary Figure 2. InferCNV analysis.**
InferCNV was utilized to infer copy number variation and determine clusters that were likely malignant versus benign.

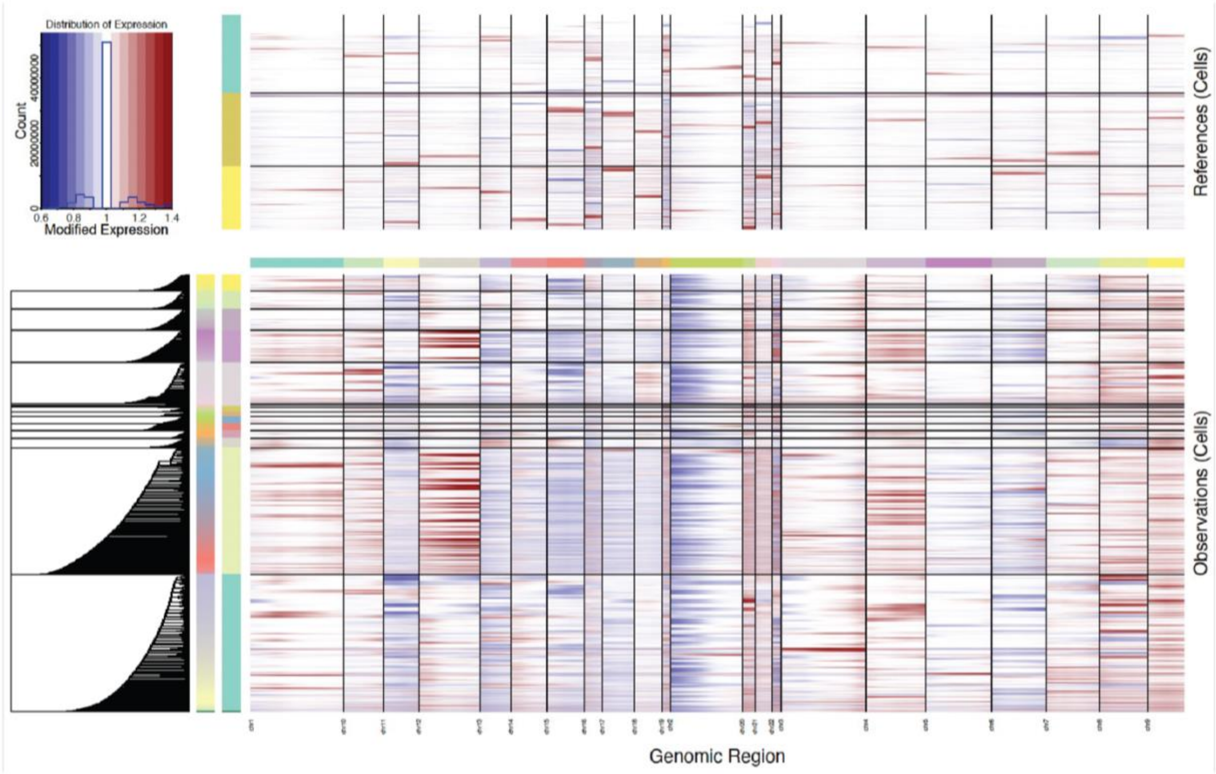

**Supplementary Figure 3. Immune microenvironment in leiomyosarcoma.**

(A) UMAP analysis of snRNA-seq data highlighting non-malignant cell types. (B) Quantification of the percent of tumor cells within each of the non-malignant clusters. Each dot represents an individual tumor, and red bars represent means ± SEM. (C) Bubble plot showing expression of general macrophage, inflammatory (classically M1), and immunosuppressive (classically M2) macrophage markers by tumor. (D) Bubble plot showing expression of markers of pan-T cells, exhausted T cells, cytotoxic T cells, and regulatory T cells (Treg). (E) Bubble plot showing expression of T cell and macrophage checkpoints in the non-malignant cell clusters.

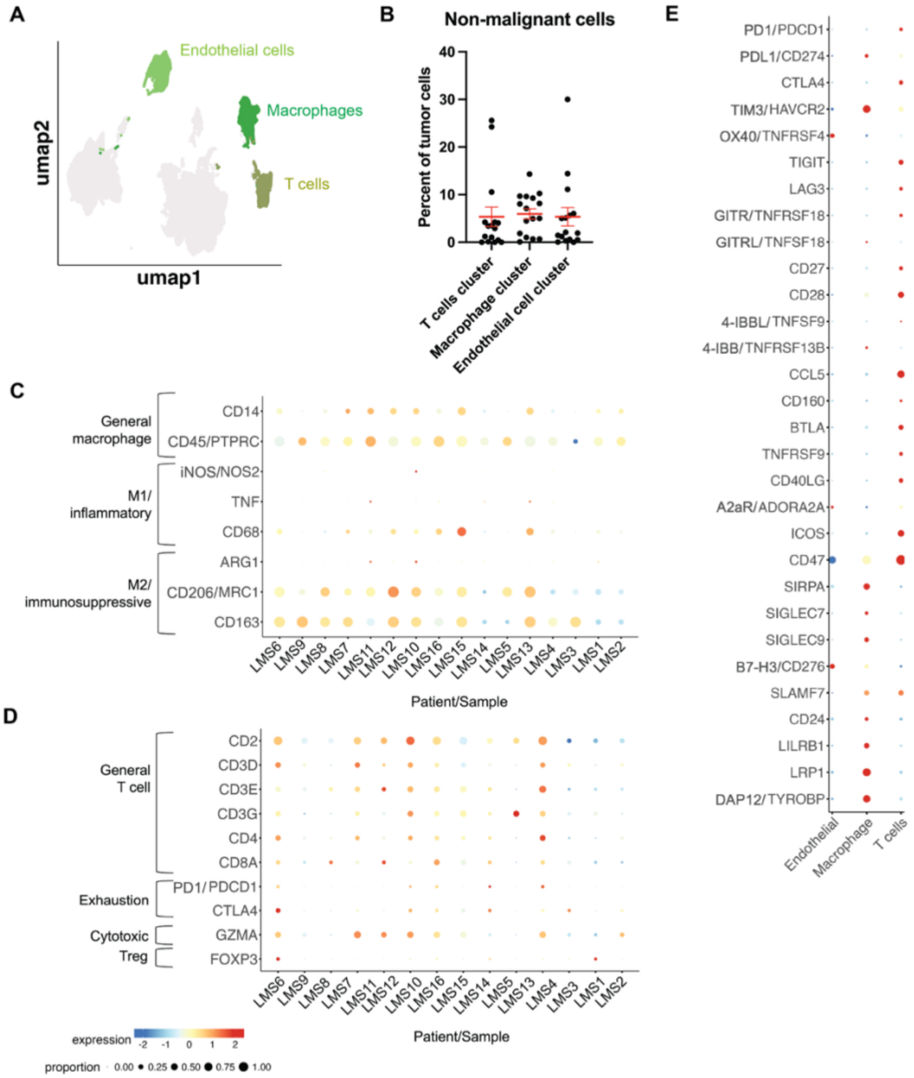

**Supplementary Figure 4. T cell subsets in leiomyosarcoma.**
Bubble plots showing expression of genes that define each T cell subset by each individual tumor.

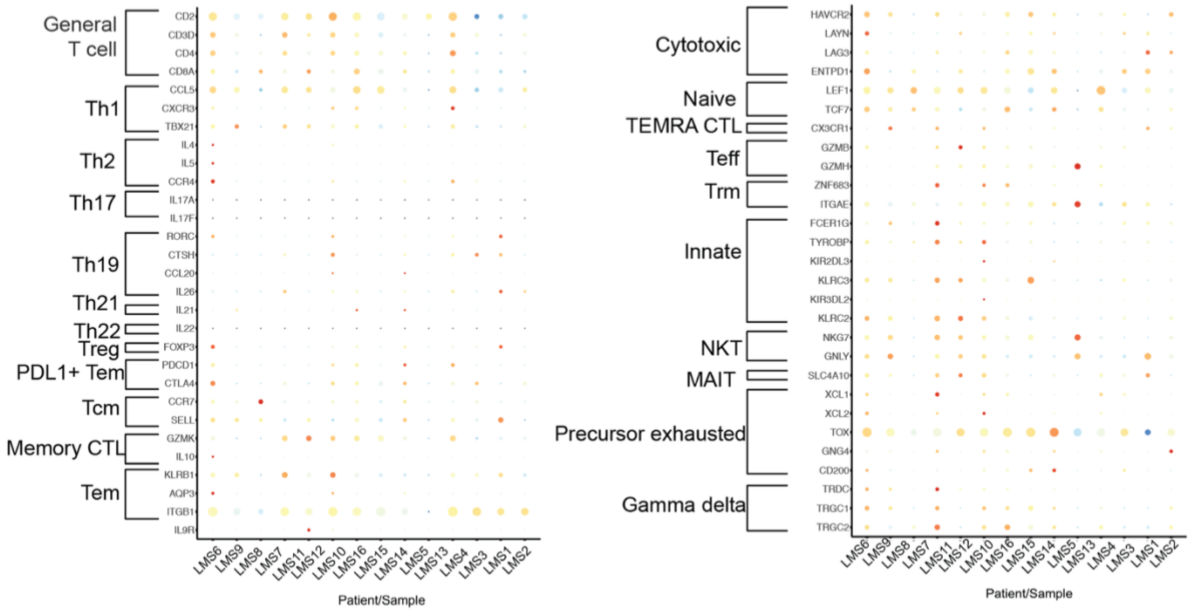

**Supplementary Figure 5. Macrophage subsets in leiomyosarcoma.**
Bubble plots showing expression of genes that define each macrophage subset by anatomic location (A) and individual tumor (B).

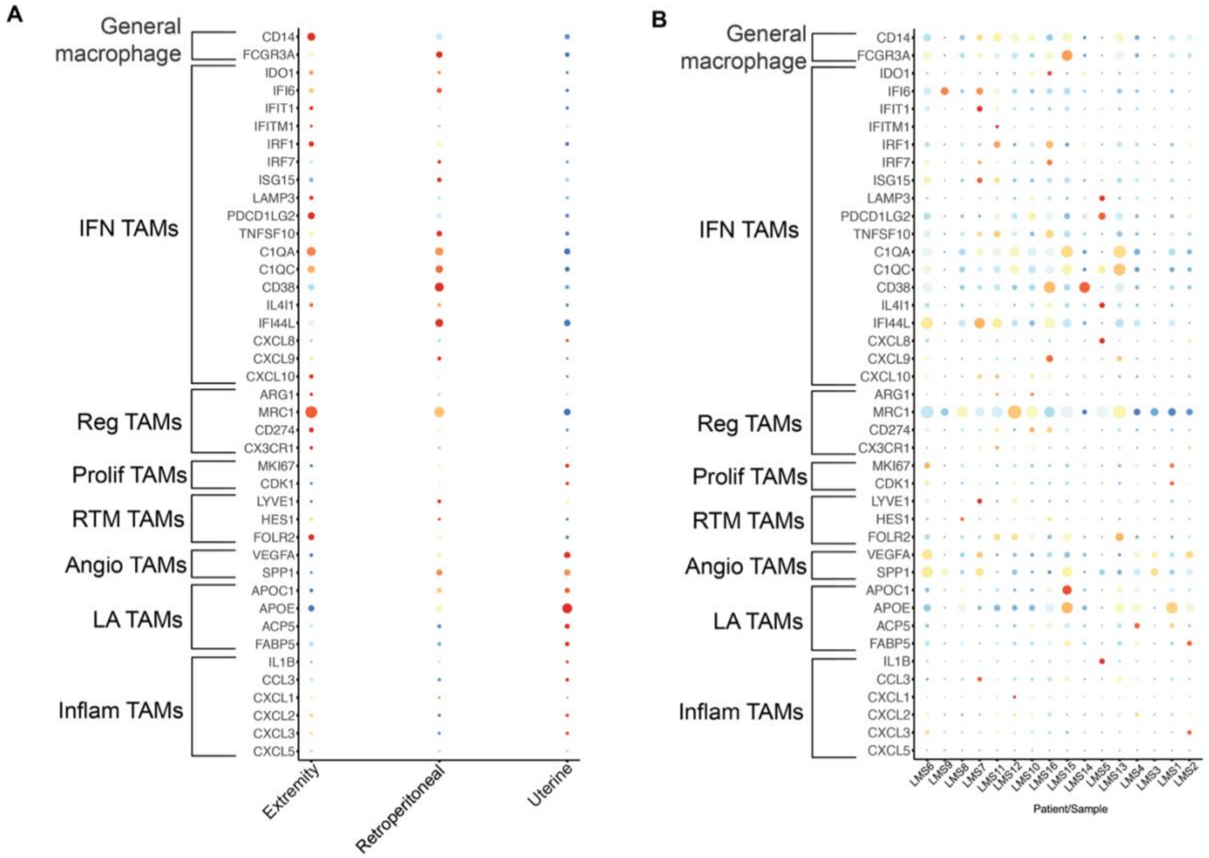

**Supplementary Figure 6. snRNA-seq analysis of 16 leiomyosarcoma tumors.**
(A) snRNA-seq UMAP by tumor/sample. (B) Proportion of cells in each tumor/sample that fell within each cluster. (C) Absolute number of cells in each tumor/sample that fell within each cluster. (D) Proportion of cells in each cluster that came from each tumor/sample. (E) Absolute number of cells in each cluster that came from each tumor/sample. (F) snRNA-seq UMAP by site of disease (retroperitoneal, extremity, uterus). (G) Proportion of cells in each site of disease that fell within each cluster. (H) Proportion of cells that fell within each cluster by site of disease.

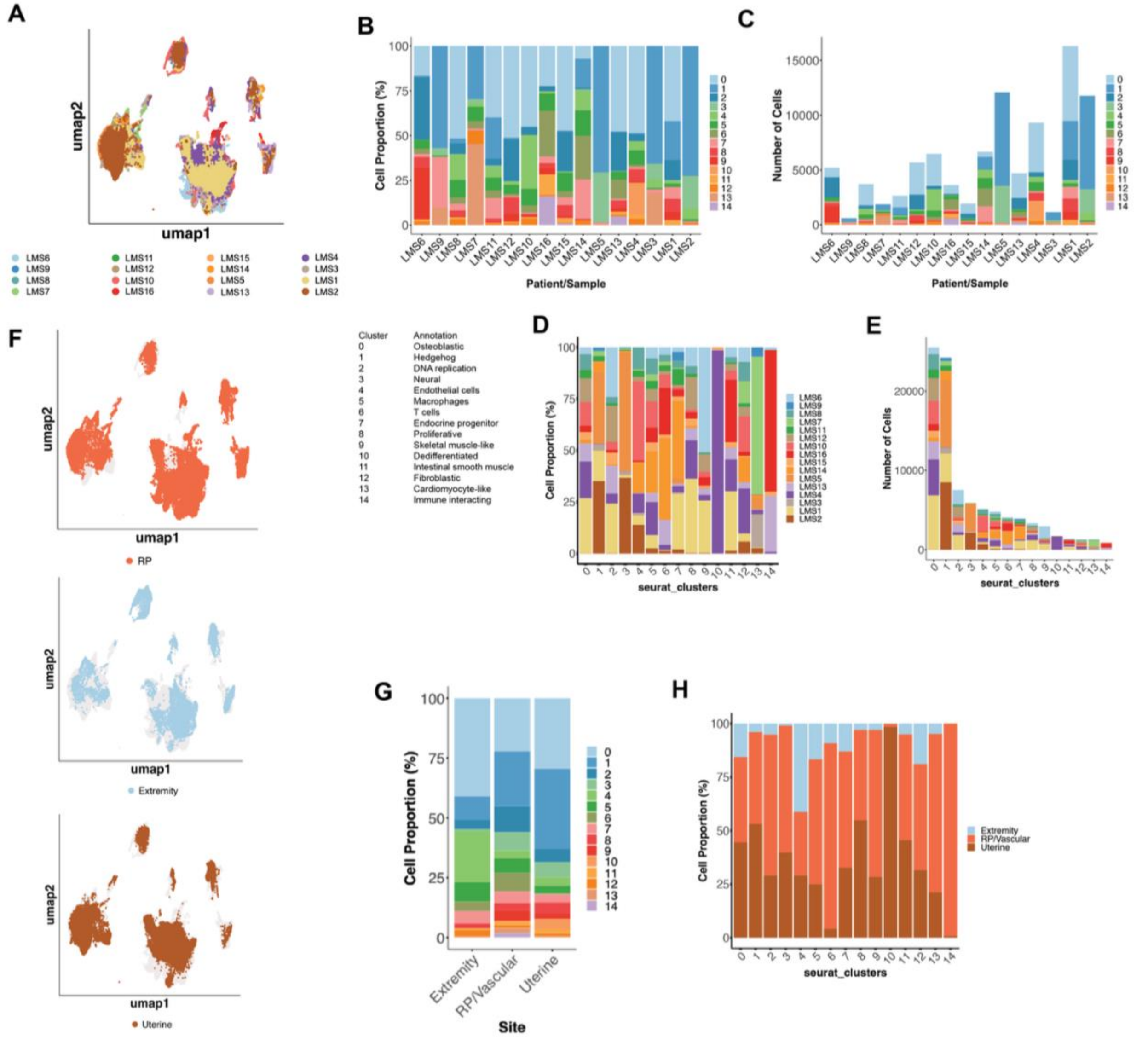

**Supplementary Figure 7. Gene set enrichment analysis by anatomic site.**
(A-C) GSEA showing hallmark pathways enriched in the differentially expressed genes for retroperitoneal leiomyosarcoma samples (A), extremity leiomyosarcoma samples (B), and uterine leiomyosarcoma samples (C). (D) Bubble plot showing enrichment of hallmark pathways in each site of disease.

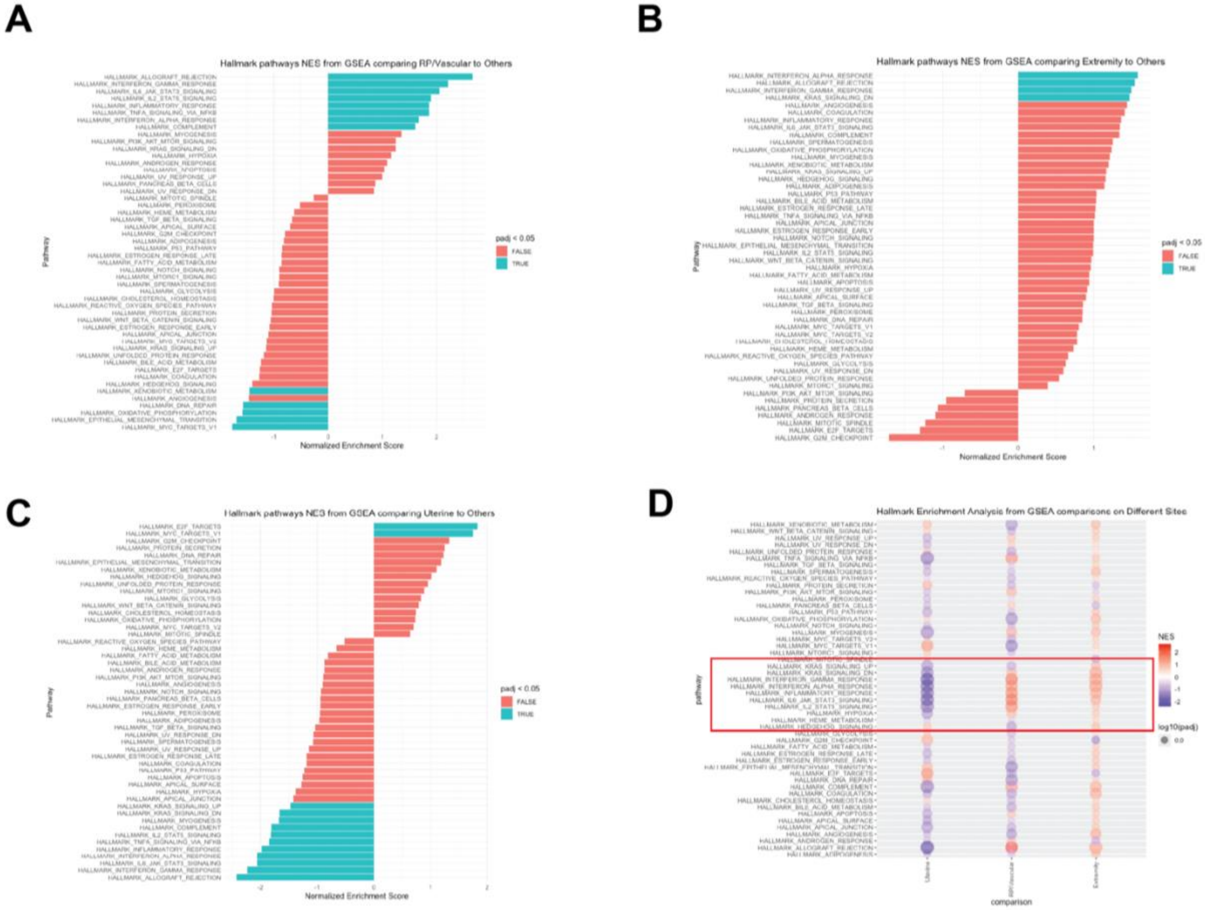

**Supplementary Figure 8. Comparisons to previously reported subtypes from bulk transcriptomic data.**

Bubble plots showing each tumor's expression of the markers that defined each of the 3 subtypes from studies by Hemming et al (A), Beck et al (B), Chudasama et al (C), and Guo et al (D).

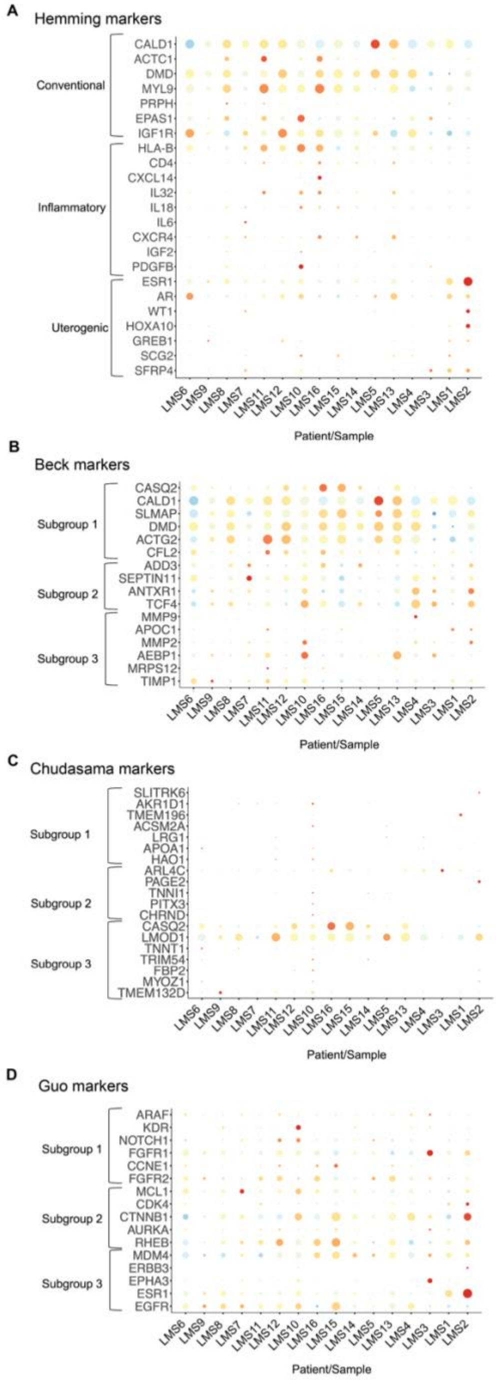

**Supplementary Figure 9. Expression of smooth muscle markers.**

(A) Bubble plot showing expression of smooth muscle markers by cluster. (B) Bubble plot showing expression of smooth muscle markers by cell subsets. (C) Bubble plot showing expression of smooth muscle markers by each individual tumor. (D) UMAPs from snRNA-seq data with each smooth muscle marker gene's expression overlaid. (E) Bubble plot showing expression of markers that define different smooth muscle phenotypes by cluster.

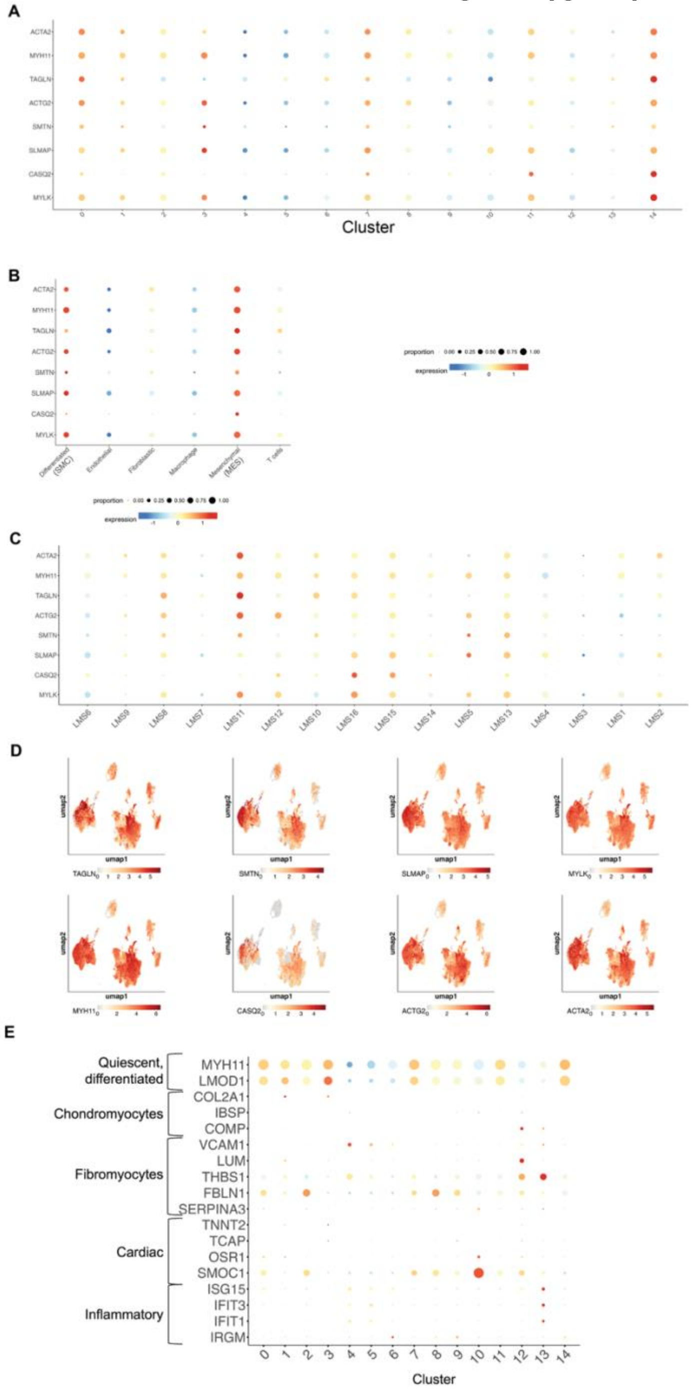

**Supplementary Figure 10. snRNA-seq UMAPs by sample.**

Each tumor's cells are projected in a different color onto the snRNA-seq UMAP. Tumors are separated by site of disease.

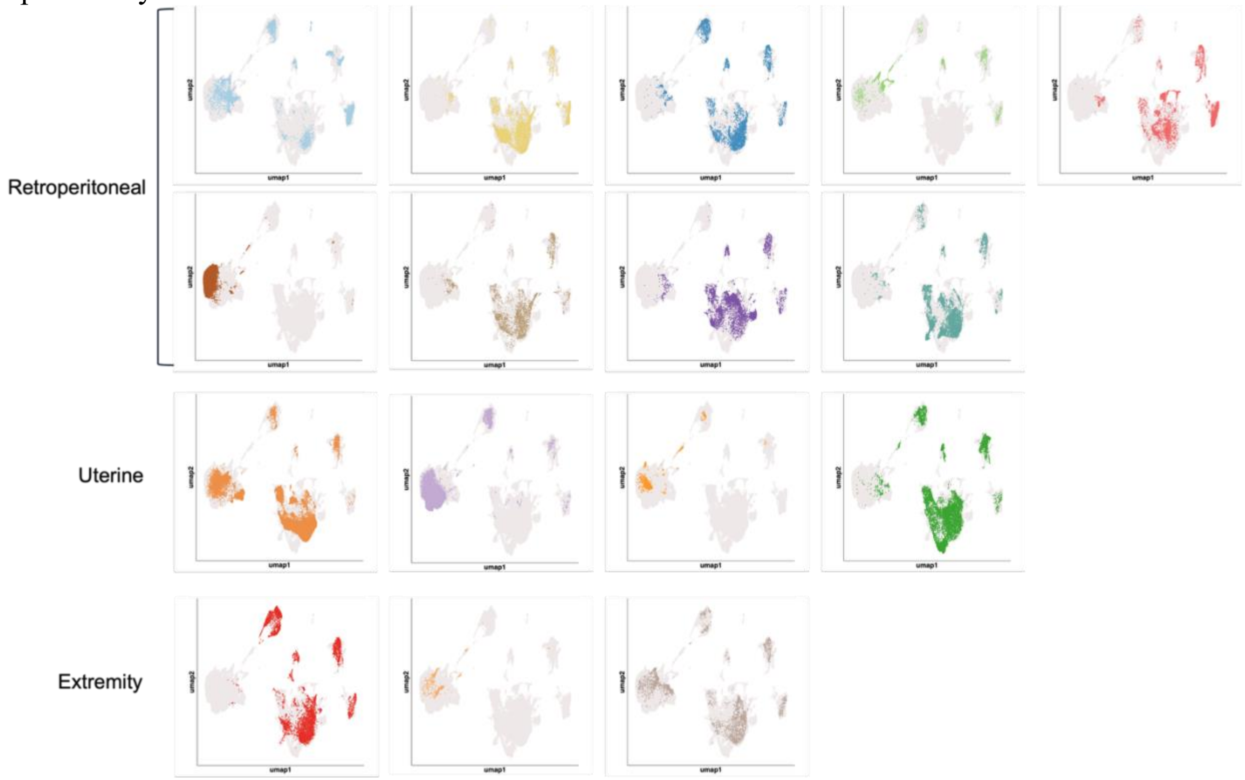

**Supplementary Figure 11. Whole genome sequencing.**

(A) Oncoplot demonstrating genomic alterations in each tumor identified by whole genome sequencing. Tumor site and grade are also shown on the top. (B-E) snRNA-seq UMAPs showing just the tumors with the indicated alteration (colored), including ATRX or DAXX loss-of-function mutation (B), DNA damage response pathway mutation (C), 17p amplification (D), and PTEN loss-of-function mutation (E).

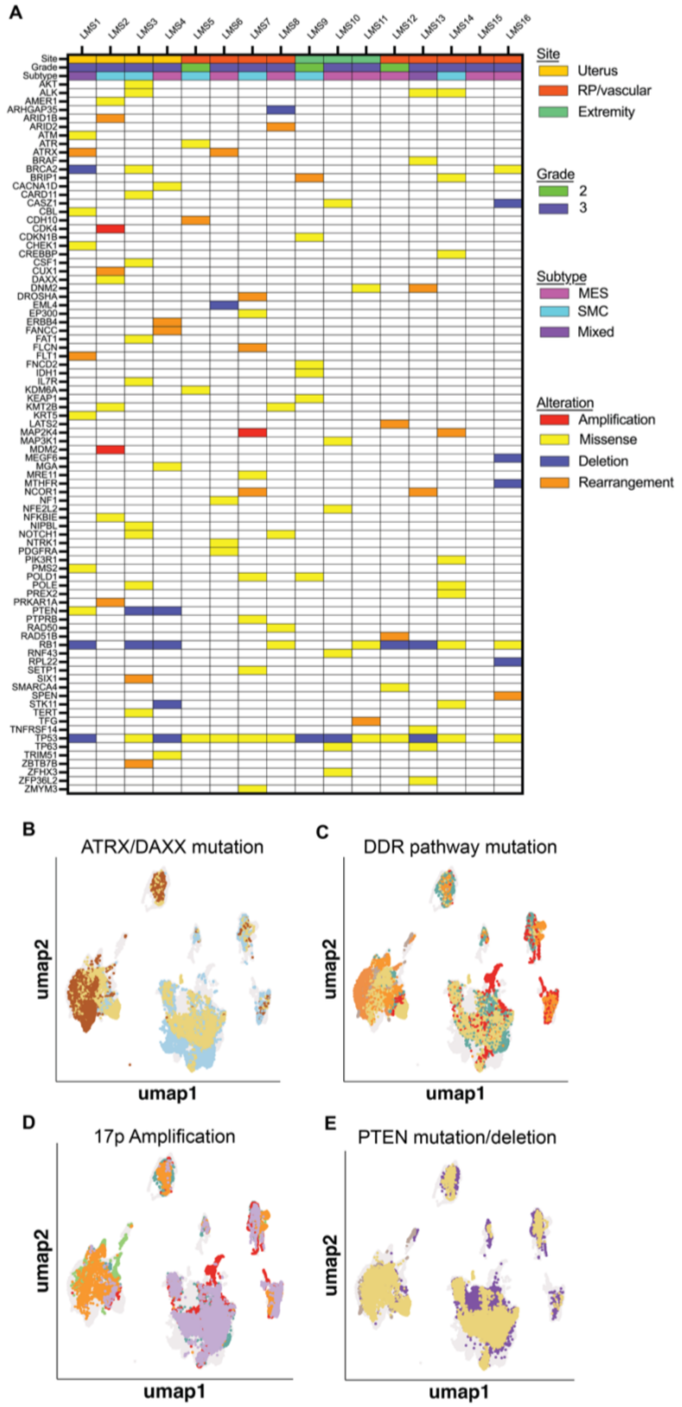

**Supplementary Figure 12. METAflux analysis.**
METAflux was used to infer metabolic flux by each cell subset.

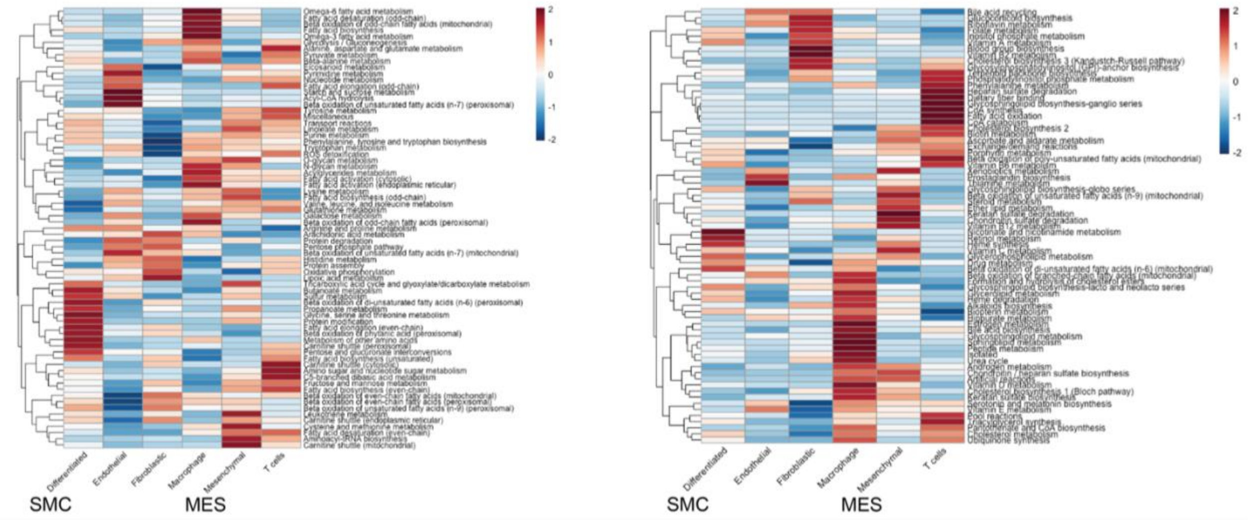

**Supplementary Figure 13. Immune microenvironment in MES- versus SMC-predominant tumors.**

(A-D) Quantification of the percent of macrophages (A), T cells (B), endothelial cells (C), and total non-malignant cells (D) within MES- and SMC-predominant tumors. (E) Bubble plot showing expression of chemokines, cytokines, and their receptors by tumor. (F) Bubble plot showing expression of chemokines, cytokines, and their receptors comparing MES to SMC subtypes.

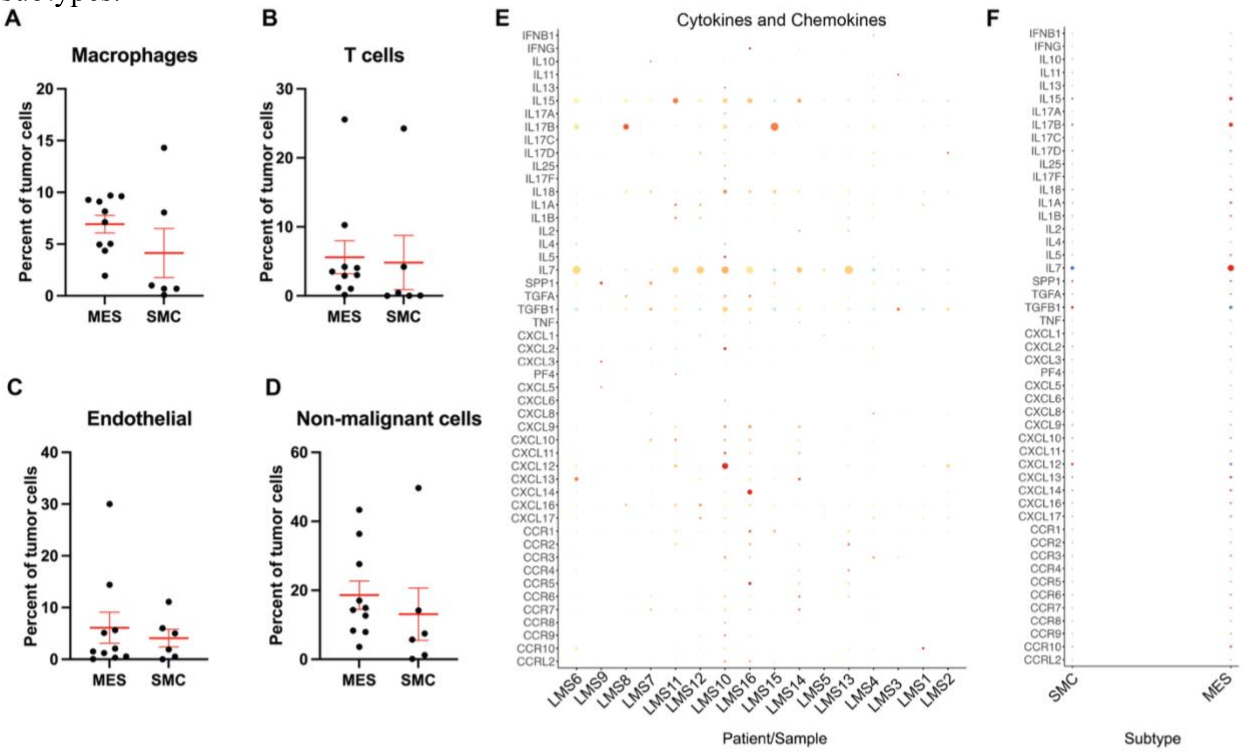

**Supplementary Figure 14. Cell interactions.**

Cell interactions were inferred by CellChat, which analyzes expression of receptor-ligand pairs. (A) Plot of the number (left) and strength (right) of interactions by cell subset. (B) Plots of interactions by each individual cell subset. (C) Heatmap showing the strength of interactions between. (D-F) The strongest receptor-ligand interactions seen involved collagen (D), laminin €, and fibronectin (F).

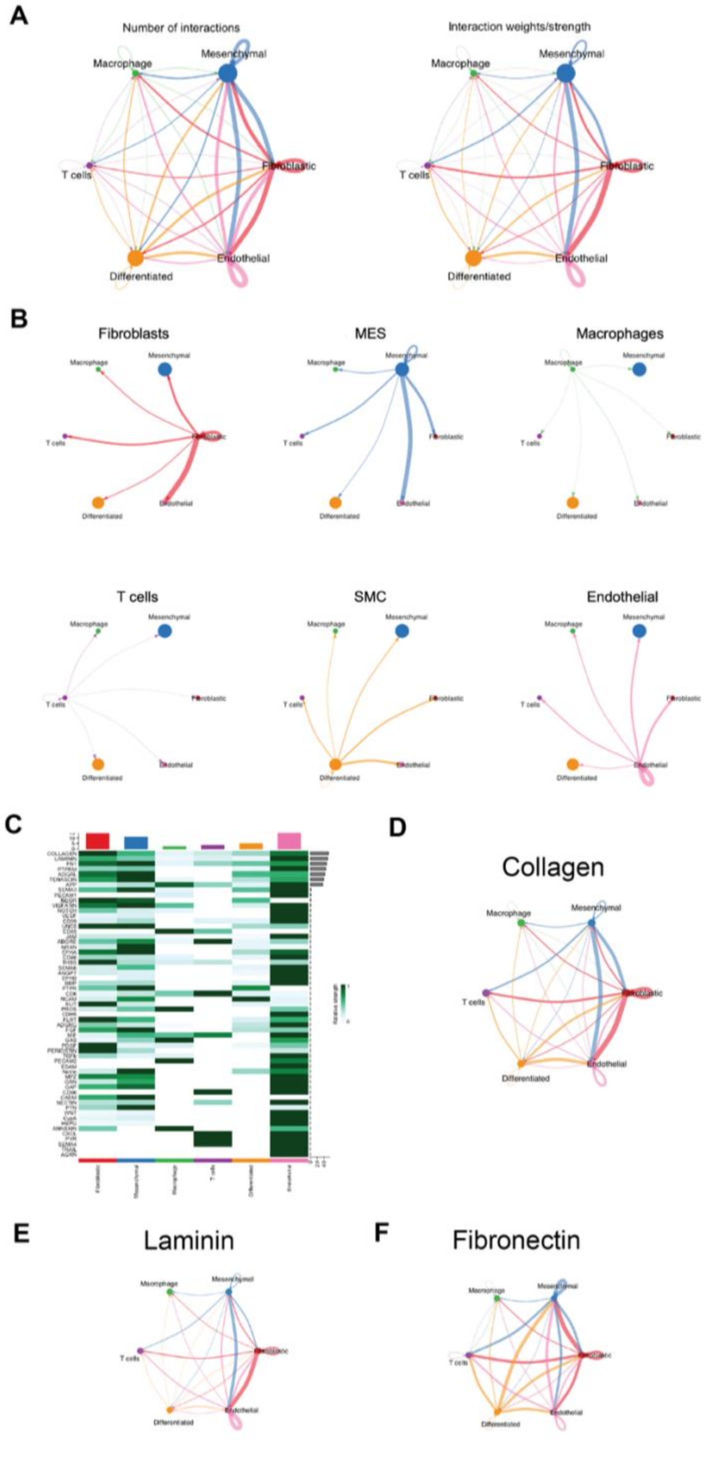

**Supplementary Figure 15. Xenium spatial transcriptomic analysis of leiomyosarcoma TMAs.**

(A-B) H&E images of leiomyosarcoma TMAs analyzed, separated by STLMS (A) and ULMS (B). (C-D) UMAP analysis of STLMS slides (C) and ULMS slides (D). (E-F) Bubble plots showing marker genes used to annotate clusters in STLMS (E) and ULMS (F). (G) Percent of leiomyosarcoma tumors by predominant subtype.

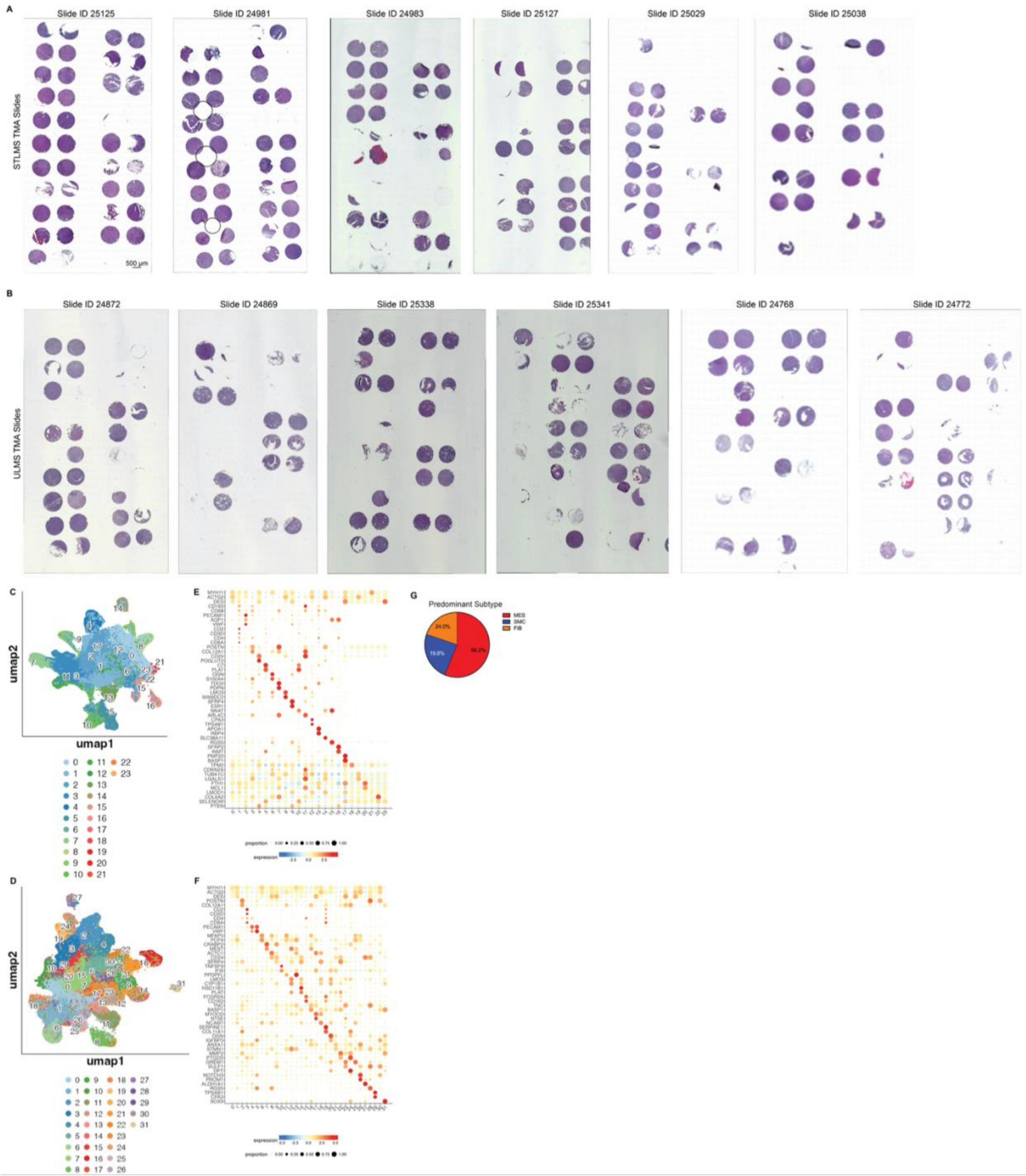

**Supplementary Figure 16. Clinical data associated with cohort of 16 patients.**
This figure shows clinical data associated with the 16 leiomyosarcoma tumors that were used for single cell multiome analysis. (A-B) Overall survival (A) and metastasis-free survival (B) comparing MES- and SMC-predominant tumors. (C-F). Percent of cells in each tumor that fall within the MES cluster are plotted based on grade (C), stage (D), site (E), and mitotic rate (F). There were no statistically significant differences in these analyses.

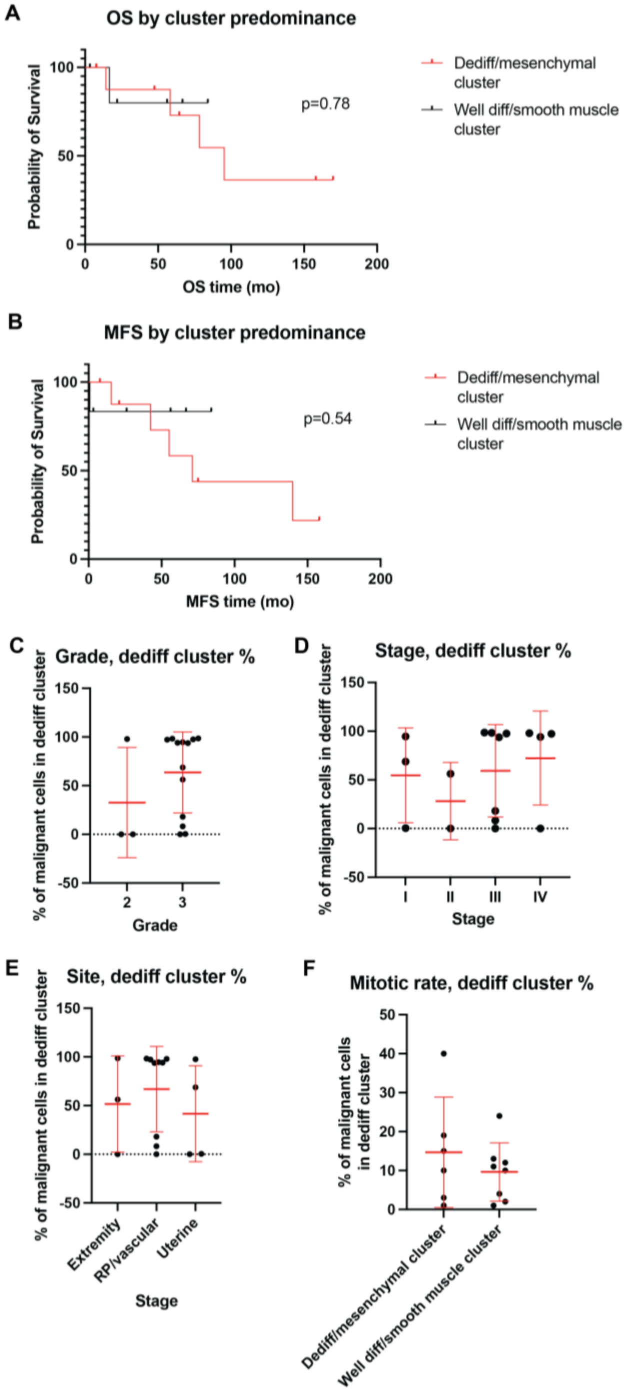

**Supplementary Figure 17. snATAC-seq analysis.**

(A) UMAP of snATAC-seq data with each color representing a different tumor. (B) UMAP of snATAC-seq data with each color representing a different cluster (from snRN-Aseq annotation). (C-D) Top enriched transcription factor (TF) motifs by subtype (C) and cluster (D). MES = mesenchymal; SMC = smooth muscle cell.

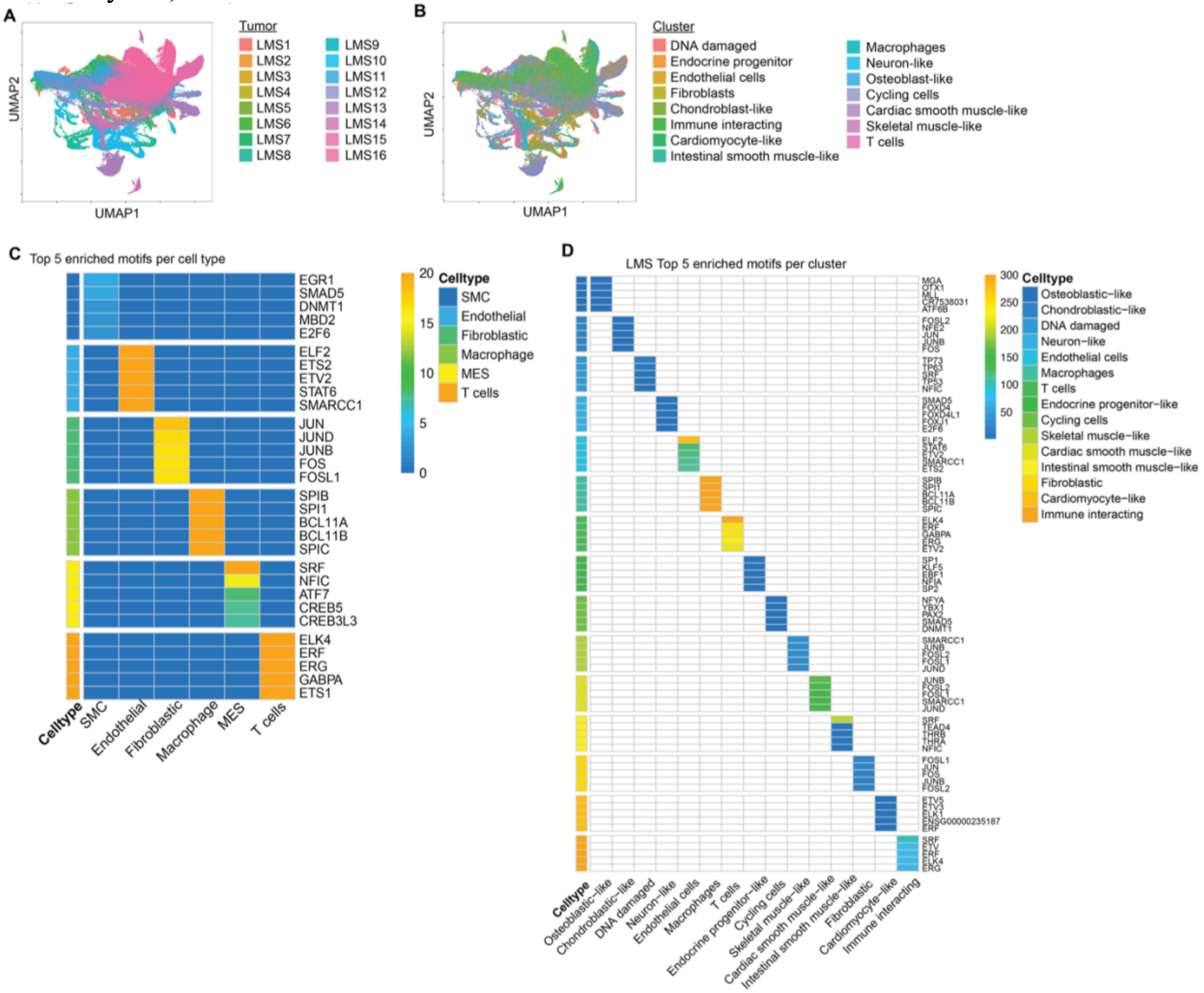

**Supplementary Figure 18. Transcription factor motif analysis.**

(A) Most upregulated TF motifs in the entire dataset when including all cells. (B) Most downregulated TF motifs in the entire dataset when including all cells. (C) Most upregulated TF motifs in the entire dataset when including only malignant cells. (D) Most downregulated TF motifs in the entire dataset when including only malignant cells. (E) Volcano plot showing TF motifs upregulated in MES versus SMC cell clusters. (F) As one tumor (LMS4) was making large contributions to this analysis, we performed a sensitivity analysis by excluding LMS4 and repeating the differential motif analysis.

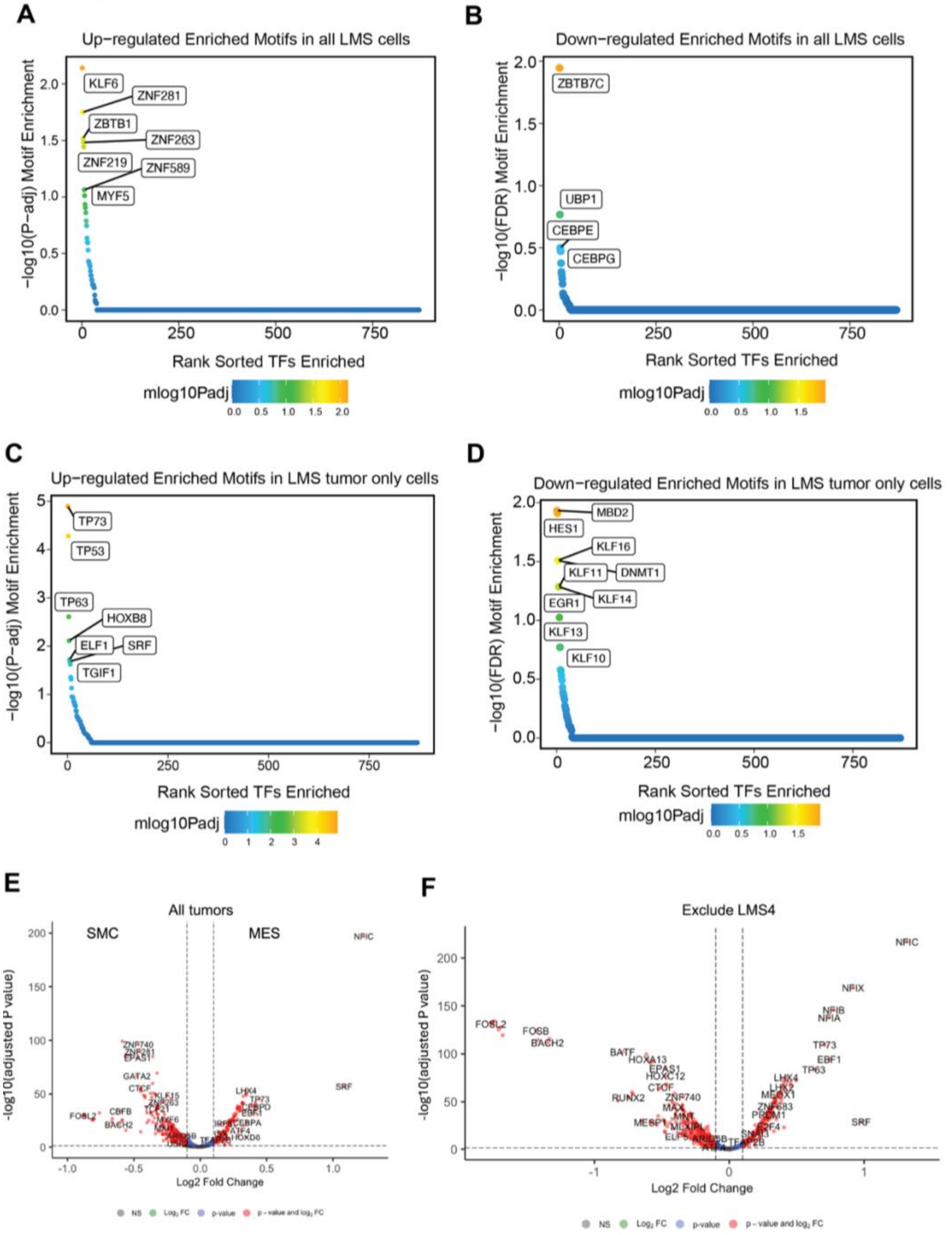

**Supplementary Figure 19. Expression of TFs in snRNA-seq data.**

(A-C) UMAPs from snRNA-seq data with each TF's gene expression overlaid for NFI TFs (A), FOS TFs (B), and JUN/BACH TFs (C). (D) Bubble plot of TF expression by cell type. (E) Bubble plot of TF expression by MES or SMC cluster. (F) Bubble plot of TF expression by MES or SMC cluster. Expression values are not scaled in this plot (but are scaled in E). (G) Plot of raw transcript count of each TF by MES or SMC cluster from snRNA-seq data.

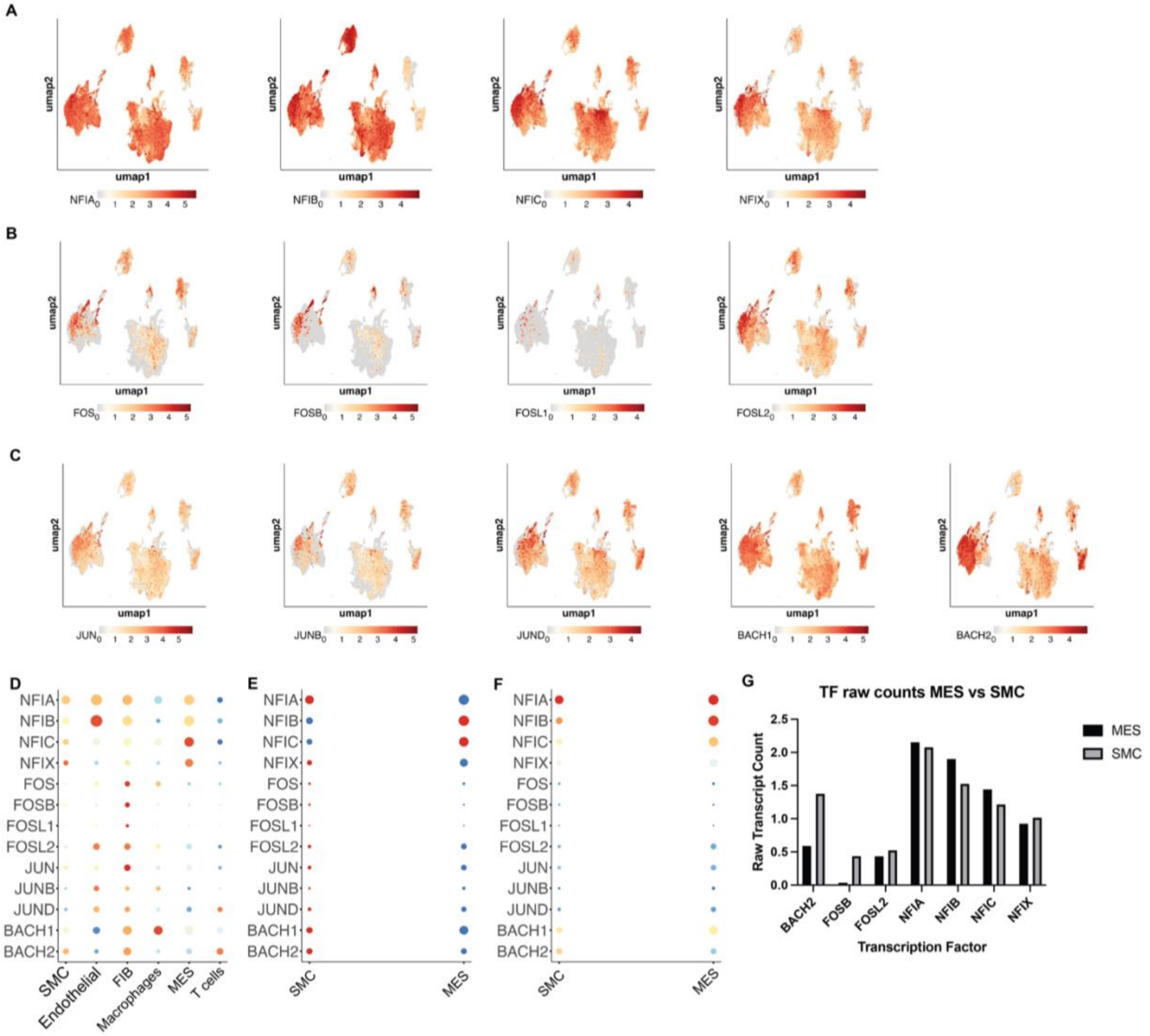

**Supplementary Figure 20. Spatial ATAC-seq.**

Spatial ATAC-seq was performed on 2 leiomyosarcoma tumors. (A) UMAP showing the 2 different leiomyosarcoma tumors in different colors. (B) UMAP showing the 10 distinct clusters that were identified. (C) Distribution of cluster by sample. (D) Top 5 genes enriched in each cluster. (E) Heat map showing the top TFs enriched by cluster. (F) UMAP showing spatial distribution of clusters. (G) UMAPs with projection of MRC1 expression to show macrophage localization. (H-K) UMAP with projection of NFIA activity (H), NFIB activity (I), NFIC activity (J), and NFIX activity (K).

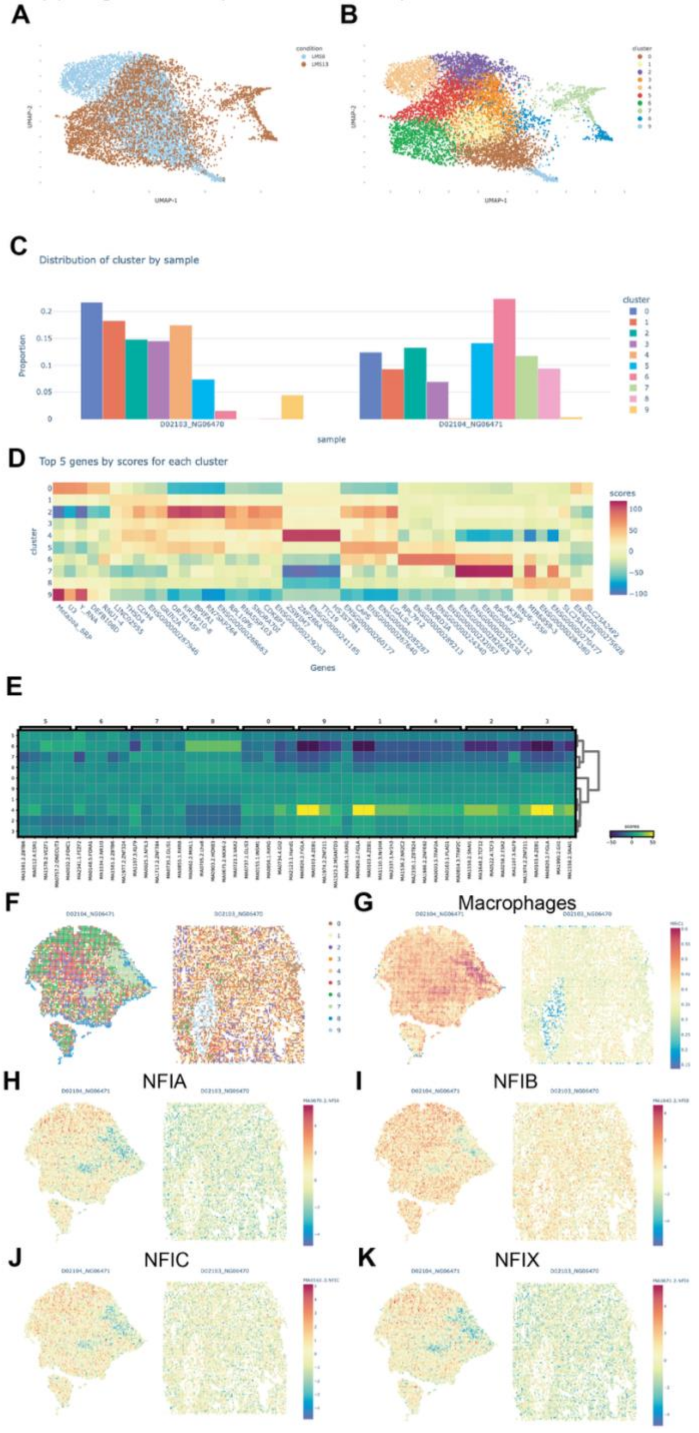

**Supplementary Figure 21. Validation of TF antibodies.**

(A) Nuclear and cytoplasmic extracts of HUtSMC and 3 leiomyosarcoma cell lines were probed for antibodies targeting each TF. Lamin B1 was used as a control to verify nuclear enrichment, and vinculin was used as a control to verify cytoplasmic enrichment. (B) Representative microphages of immunofluorescent staining using antibodies targeting the indicated TFs. Scale bar = 100 μm.

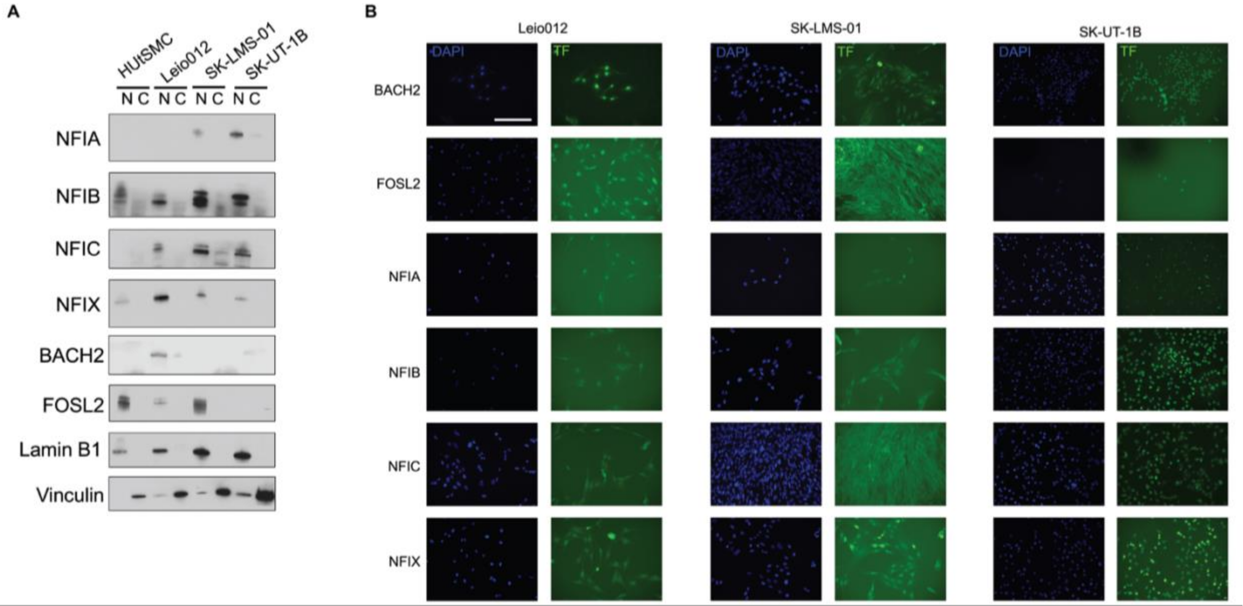

**Supplementary Figure 22. Depletion of TFs in leiomyosarcoma cell lines.**
(A) Western blots showing depletion of the indicated TFs by shRNAs. (B) Depletion of AP-1 TFs from SK-UT-1B (MES phenotype), which does not express these TFs. (C) Depletion of NFI TFs from Leio-012 (SMC phenotype), which does not express these TFs.

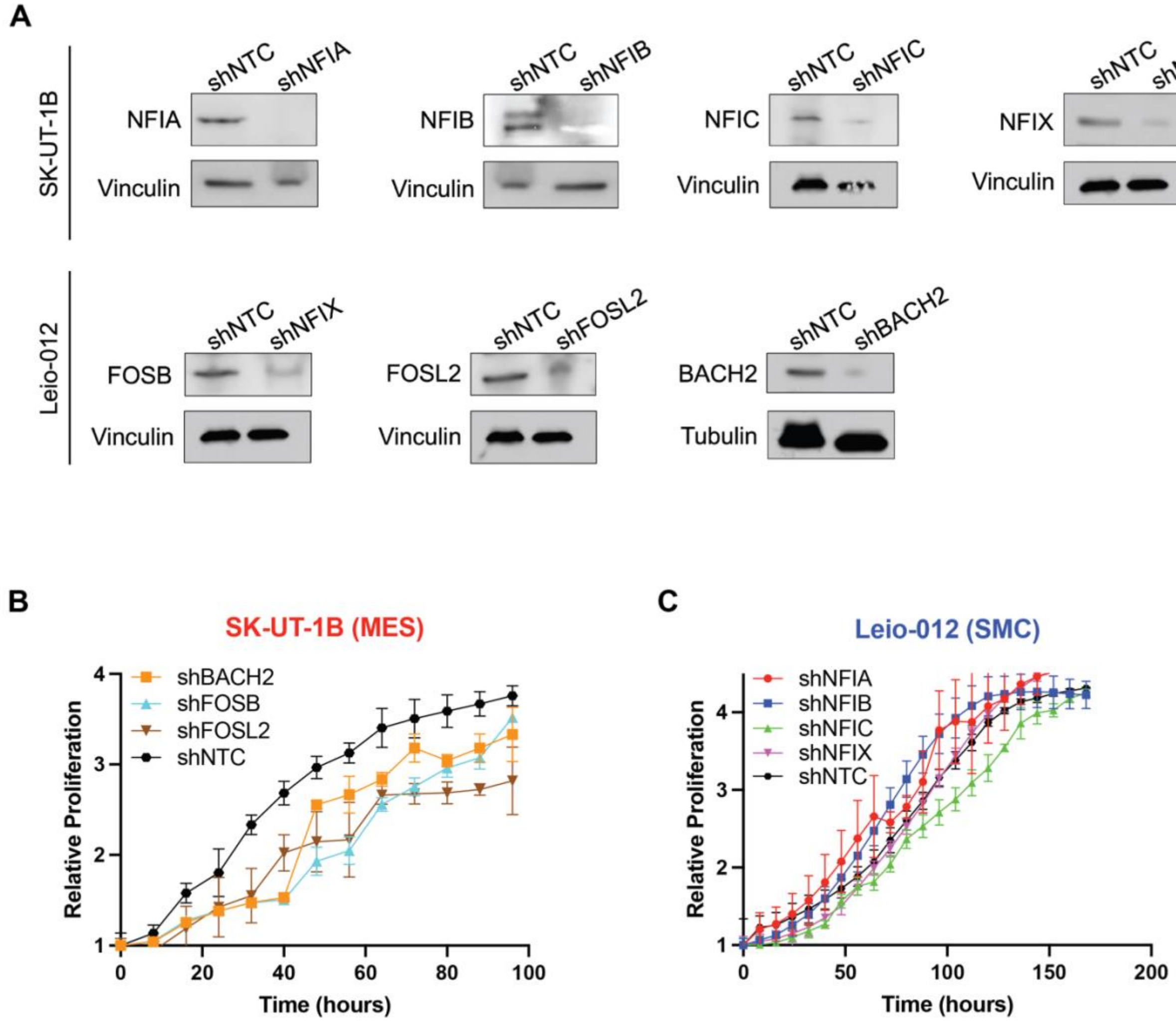

**Supplementary Figure 23. Trajectory analysis.**
Trajectory velocities determined by scVelo projected onto the snRNA-seq UMAP.

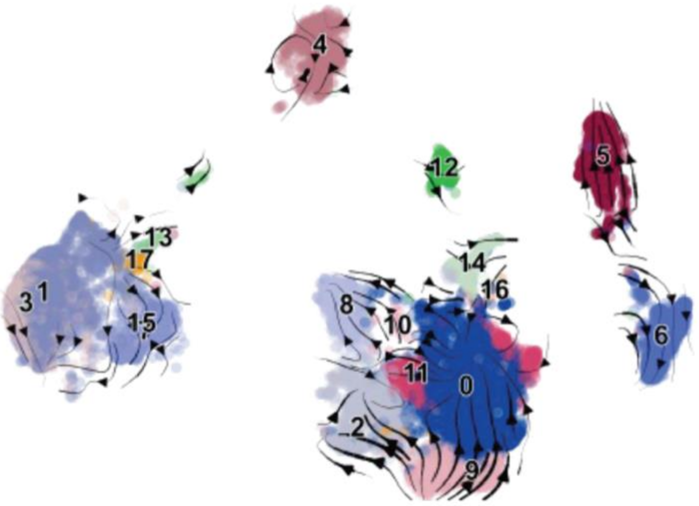

**Supplementary Figure 24. Transcription factor chromatin immunoprecipitation.**
(A-B) Plots demonstrating distribution of the peaks identified by ChIP for each indicated transcription factor (TF) throughout the genome for SK-UT-1B (A) and Leio-012 (B-D). (C) Pathway analysis using Enrichr using the genes whose peaks were identified as targets of the indicated TFs from ChIP in SK-UT-1B (C) and Leio-012 (D).

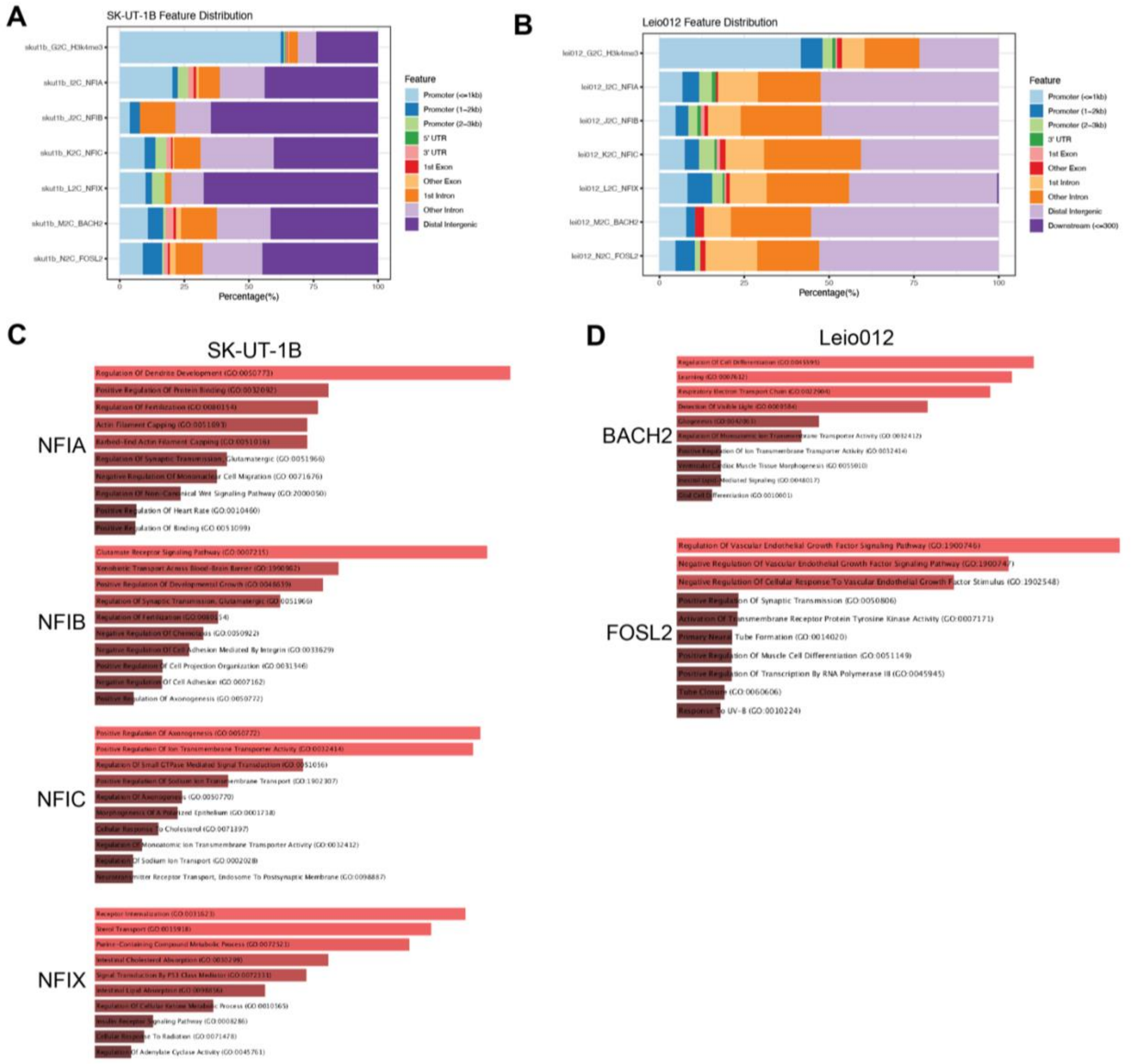

**Supplementary Figure 25. Epigenetic drug screen.**
Leiomyosarcoma cell lines were screened with a panel of epigenetic drugs representing the classes of agents that are currently available and either FDA-approved or in clinical trials.

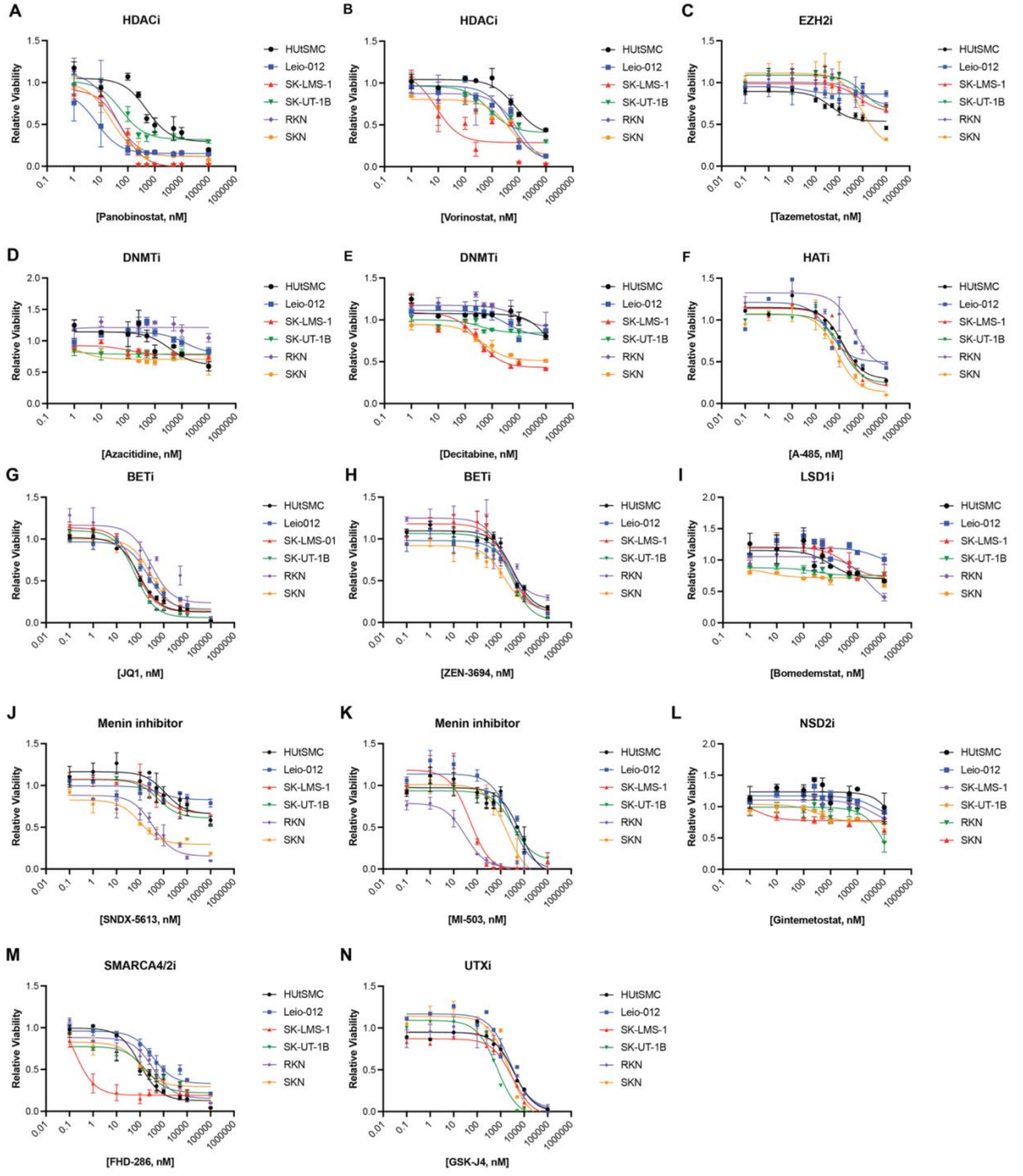

**Supplementary Figure 26. Apoptosis assay gating strategy.**

Leiomyosarcoma cell lines were treated with FHD-286 and analyzed for apoptosis using Annexin V FITC and propidium iodide (PI) staining. This figure demonstrates the gating strategy used, filtering all cells (upper left), gating on single cells (upper right), and dividing cells into living, early apoptosis, late apoptosis, and necrosis based on Annexin V FITC and PI staining.

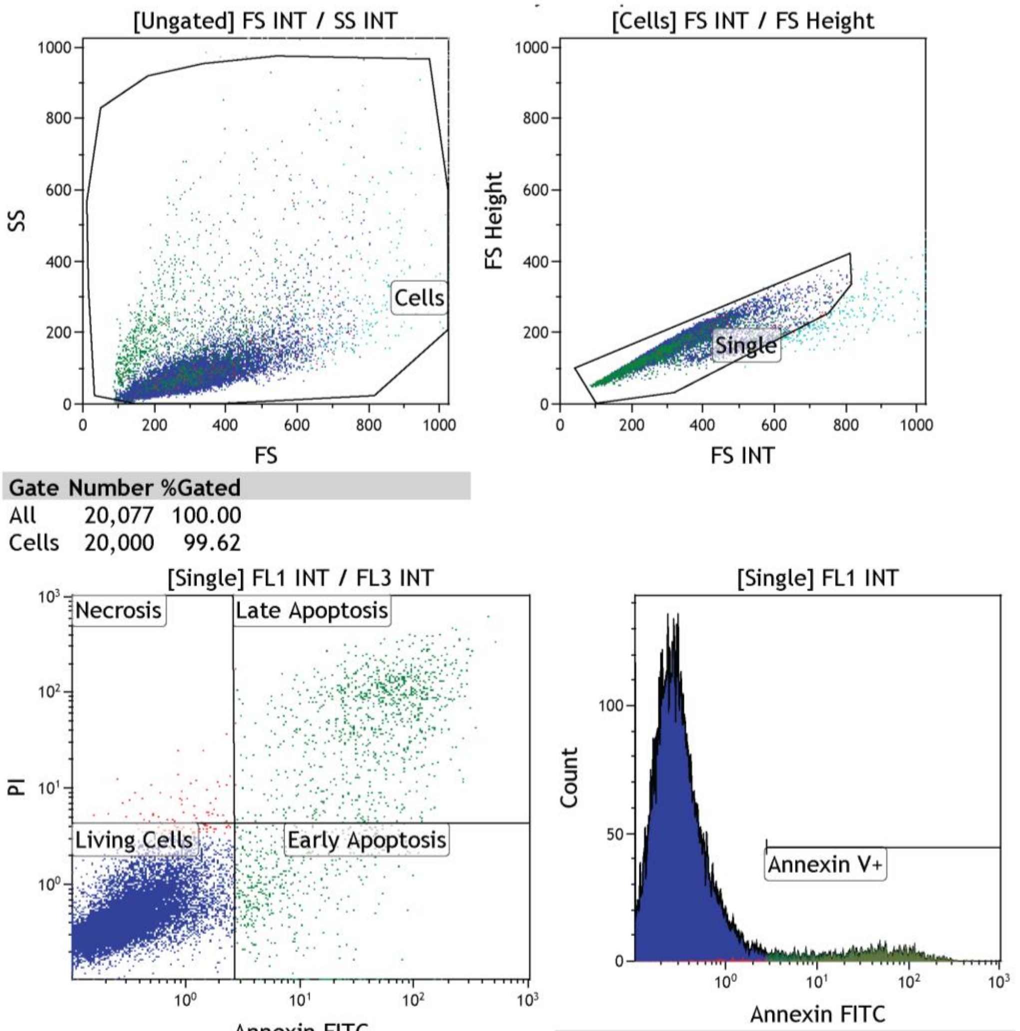


| Gate | Number | %Gated |
|---|---|---|
| All | 20,077 | 100.00 |
| Cells | 20,000 | 99.62 |

| Gate | Number | %Gated | X-Med | Y-Med |
|---|---|---|---|---|
| All | 19,762 | 100.00 | 0.30 | 0.44 |
| Early Apoptosis | 513 | 2.60 | 7.49 | 1.04 |
| Late Apoptosis | 936 | 4.74 | 50.02 | 72.10 |
| Living Cells | 18,257 | 92.38 | 0.28 | 0.42 |
| Necrosis | 56 | 0.28 | 1.03 | 6.08 |

| Gate | Number | %Gated |
|---|---|---|
| All | 19,762 | 100.00 |
| Annexin V+ | 1,425 | 7.21 |
